# BRG1/BRM inhibitor targets AML stem cells and exerts superior preclinical efficacy combined with BET or Menin inhibitor

**DOI:** 10.1101/2023.09.28.560054

**Authors:** Warren Fiskus, Jessica Piel, Mike Collins, Murphy Hentemann, Branko Cuglievan, Christopher P. Mill, Christine E. Birdwell, Kaberi Das, John A. Davis, Hanxi Hou, Antrix Jain, Anna Malovannaya, Tapan M. Kadia, Naval Daver, Koji Sasaki, Koichi Takahashi, Danielle Hammond, Patrick Reville, Jian Wang, Sanam Loghavi, Rwik Sen, Xinjia Ruan, Xiaoping Su, Lauren B. Flores, Courtney D. DiNardo, Kapil N. Bhalla

## Abstract

BRG1 (SMARCA4) and BRM (SMARCA2) are the core ATPases of chromatin remodeling BAF (BRG1/BRM-associated factor) complexes, which enable transcription factors/co-factors to modulate gene-expressions, mediating growth, differentiation-arrest and survival of AML cells. In AML with MLL1r (MLL1 rearrangement) or mutant (mt) NPM1, although monotherapy with Menin inhibitor (MI) induces clinical remissions, most patients either fail to respond or relapse. FHD-286 is a selective BRG1/BRM inhibitor, undergoing clinical development in AML. Here, FHD-286 induced differentiation and lethality in AML cells with MLL1r or mtNPM1, reducing chromatin accessibility and repressing c-Myc, PU.1 and CDK4/6. Whereas FHD-286 monotherapy reduced AML burden, leukemia-initiating potential and improved survival, FHD-286 combinations with MI, BET inhibitor, decitabine or venetoclax was significantly more effective in reducing AML burden and improved survival, without significant toxicity, in xenograft models of AML with MLL1r or mtNPM1. These findings highlight promising FHD-286-based combinations for therapy of AML with MLL1r or mtNPM1.

**Statement of Significance:** Inhibition of BRG1/BRM ATPases by FHD-286 reduced chromatin accessibility, repressed c-Myc, PU.1 and CDK4/6, inducing differentiation, leukemia-initiating potential and lethality in AML stem-progenitor cells. FHD-286-based combinations with Menin or BET inhibitor or decitabine reduced AML burden and improved survival in xenograft models of AML with MLL rearrangement or mutant NPM1.

## Introduction

All chromatin dynamics related to nucleosomes involve the activity of ATP dependent chromatin-modifying and remodeling complexes, which bind and allow transcription factors (TFs) and co-factors to gain access and modulate transcription (1-3). The chromatin remodeling canonical, polybromo and non-canonical BAF complexes contain mutually exclusive core ATPases, BRG1 (SMARCA4) and BRM (SMARCA2), which are composed of between 10 to 15 other protein subunits (1-4). The canonical BAF (BRG1/BRM-associated factor) complex is essential for lineage specific gene expression by TFs and for hematopoiesis (5). AML cells express and depend on BRG1/BRM (6,7). Although common in solid tumors, mutations in BRG1 or the other subunits of BAF complexes are uncommon in AML (2,6,7). Cancer cells with reduced BRG1 levels or BRG1 mutation depend for survival on BRM activity in the BAF complex (2,8). BRM depletion was shown to selectively inhibit in vitro and in vivo growth of BRG1 mutant cancer cells (2,8). Small molecule inhibitors of the ATPase activity of BRG1 and BRM have been developed, which repress BRG1/BRM-dependent gene-expression, induce differentiation and inhibit in vitro and in vivo growth of solid tumor and AML cells (8-10).

FHD286 (Foghorn Therapeutics) is a small molecule, orally bioavailable, BRG1 and BRM-selective, ATPase inhibitor. Based on its potent pre-clinical activity against cancer and leukemia cells, FHD286 is currently being evaluated for safety and clinical efficacy in early clinical trials in AML (NCT04891757). In the present studies, we interrogated the in vitro and in vivo efficacy of FHD-286, as well as its molecular correlates in models of AML cell lines and patient-derived AML cells with MLL1r or mutant (mt) NPM1, which constitute almost 40% of adult AML (11,12). This approach is supported by observations that BRG1 and BAF complex activity plays a role in the maintenance of AML with MLL1r (13). Findings presented demonstrate that treatment with FHD-286 overcame differentiation block and significantly induced in vitro differentiation and loss of viability in AML cell lines and PD AML cells with MLL1r or mtNPM1 similar to those reported earlier (14,15). FHD-286-induced lethality was associated with marked perturbations in chromatin accessibility, inhibition of enhancers, core regulatory circuitry and gene expressions in the AML cells (2,7,16,17). Our findings also demonstrate in vivo efficacy of monotherapy with FHD-286, including depletion of leukemia-initiating AML stem-progenitor cells, reduction of AML burden and significant survival gains in PDX models of AML with MLL1r or mtNPM1 (14,15,18). Whereas bulk AML reduction and achieving complete remissions is common with standard regimens, subsequent relapse and therapy refractoriness emerges in most AML patients due to persistence and enrichment of the residual AML-initiating stem-progenitor cells harboring epigenetic/adaptive escape mechanisms from standard or targeted anti-AML therapies (15,19,20). Based on the activity of FHD-286 monotherapy against AML-initiating stem-progenitor cells noted above, we also determined in vitro and in vivo efficacy of FHD-286-based combinations with standard anti-AML drugs, e.g., decitabine and venetoclax (21), as well as with promising new agents, e.g., BET inhibitor (BETi) or Menin inhibitor (MI) previously shown to exhibit pre-clinical and clinical activity in AML with MLL1r or mtNPM1 (15,22-24). Taken together, our findings below highlight the promise of FHD-286 treatment alone and in rational combinations in exerting significant anti-AML efficacy against cellular models of AML with MLL1-r or mtNPM1.

## Results

### AML cells with MLL1r or mtNPM1 are dependent on BRG1 (SMARCA4) for survival and sensitive to FHD-286-induced differentiation and loss of viability

We first interrogated the CRISPR-gRNA and RNAi dependency-screens (DepMap) to determine whether SMARCA4 is a dependency in AML cell lines (15, 25). As shown in **Fig. S1A** and **S1B**, the SMARCA4 gene-effect scores were between 0 and -1.0, highlighting it to be a dependency in several AML cell lines, including those that express MLL-fusion protein (FP) (e.g., MOLM13, MV4-11 and NOMO1) or mtNPM1 (e.g., OCI-AML3). Immunoblot analyses on cell lysates of AML cells from 16 patients, harboring either MLL1r (eight) or mtNPM1 (eight), confirmed that multiple isoforms of BRG1 and BRM proteins are expressed, albeit at disparate levels, in these PD AML cells (**Fig. S1C**). Next, we determined the anti-AML efficacy of FHD-286, a clinical grade, orally bioavailable, catalytically active-site, dual BRG1 and BRM inhibitor (10), currently undergoing Phase I clinical evaluation in patients with hematologic malignancies including AML (NCT04891757). We determined that exposure to relatively low concentrations of FHD-286 (10 to 30 nM for 7 days) induces CD11b expression, with increased mean fluorescence intensity (MFI) of CD11b, and morphologic features of differentiation, i.e., increase in % of myelocytes or metamyelocytes (15), as well as induces loss of viability in AML cells with MLL1r (MV4-11 and MOLM13) or mtNPM1 (OCI-AML3) (**Figs. 1A to 1C** and **Figs. S1D** and **S1E**). After 7 days of exposure to FHD-286, the CD11b-sorted, PD AML cells, were subsequently cultured for 7 days in drug-free medium. They showed an increased level of differentiation (**Fig. S1F**). Treatment with FHD-286 at higher concentrations (up to 100 nM) for 72 to 96 hours dose-dependently induced loss of viability in PD AML cells with MLL1r or mtNPM1 (**Fig. 1D**). In contrast, normal CD34+ progenitor cells were markedly less sensitive to similar exposures to FHD-286 (**Fig. S1G**). The oncoplot of NGS-determined co-mutations in the PD AML samples utilized in these studies is shown in **Fig. S1H**. From a study involving a larger cohort of AML samples, we had previously reported that approximately 9% of AML with MLL1-r also exhibit co-mutations in TP53, which is known to confer therapy resistance and poor outcome in AML (14, 26-28). Treatment with FHD-286 induced the same level of loss of viability in the isogenic MOLM13 cells with CRISPR knock-in of missense mutant TP53R175H or TP53R248Q, as compared to MOLM13 cells with two copies of wild-type TP53 (29) (**Fig. S1I**). After repeated exposures to LD_90_ concentrations of MI (SNDX-5613) for 96 hours, followed by recovery and growth, we generated the MI-tolerant/resistant (MITR) MV4-11 and OCI-AML3 cells (MV4-11/MITR and OCI-AML3/MITR cells) that were highly resistant to the MI SNDX-50469 (**Figs. S1J and S1K**). A qPCR assay developed to identify presence of the six, recently reported hotspot mutations in Menin that confer resistance to the activity of SNDX-50469, and its clinical grade analog SNDX-5613, failed to demonstrate any of the Menin mutations (**Fig. S1L**)(30). This suggests that resistance to MI in MV4-11/MITR and OCI-AML3/MITR cells is non-genetic and likely due to undefined adaptive/epigenetic mechanisms (31). Importantly, exposure to FHD-286 dose-dependently induced a greater level of differentiation in MV4-11/MITR and OCI-AML3/MITR cells, as compared to MV4-11 and OCI-AML3 cells (**Figs. 1E to 1H**). Together, these findings indicate that FHD-286 retains activity against AML cells with MLL1r and FLT3-ITD (MV4-11 and MOLM13 cells) or harboring TP53 mutations, as well as against MI-resistant AML cells with MLL1r or mtNPM1.

**Figure 1.**
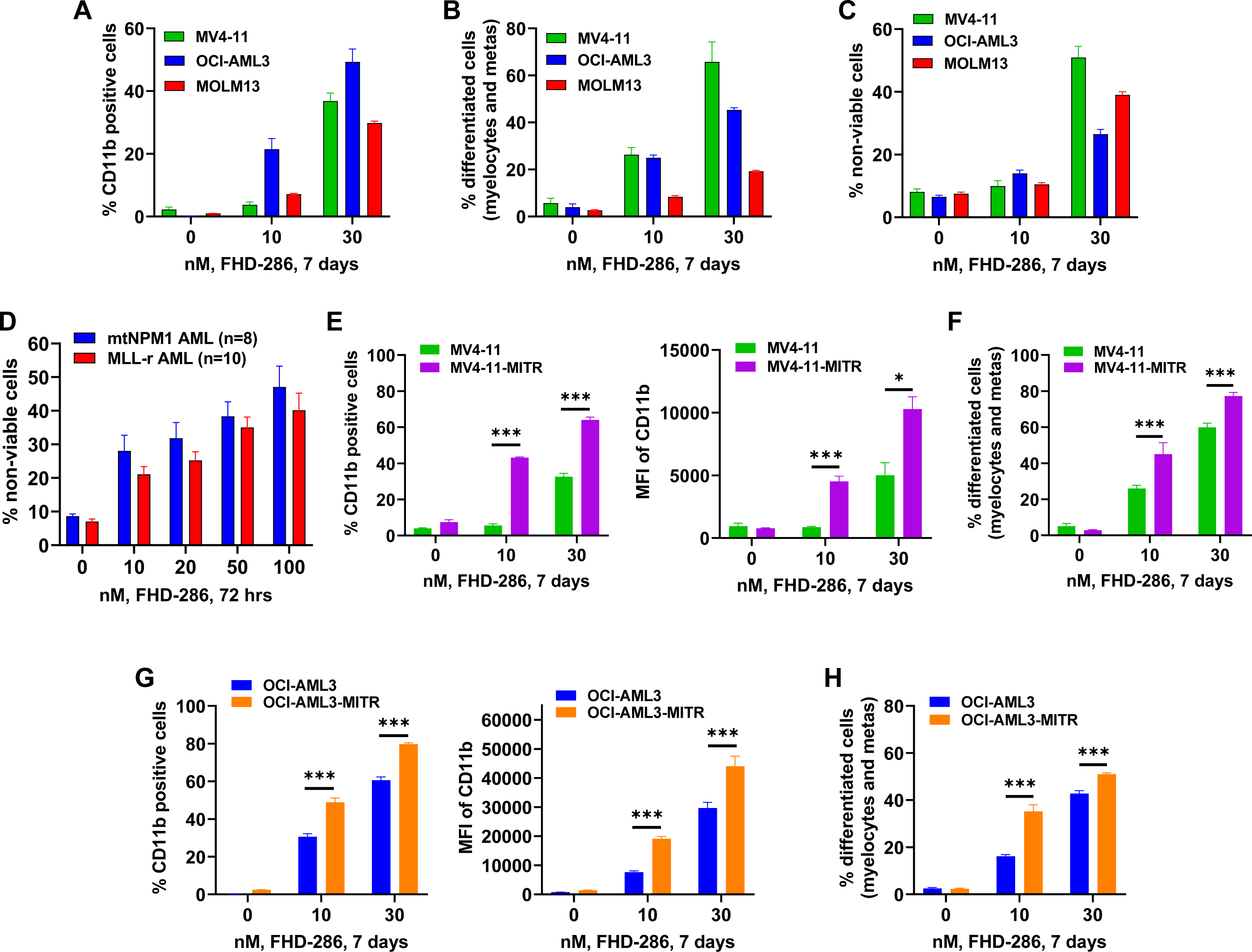
Treatment with FHD-286 overcomes differentiation block and significantly induced CD11b expression and morphologic features of differentiation in AML cell lines with MLL1r and mtNPM1. **A-C.** MV4-11, OCI-AML3 and MOLM13 cells were treated with the indicated concentrations of FHD-286 for 7 days. At the end of treatment, cells were assessed for CD11b expression, morphologic features of differentiation and percentage of non-viable cells. Columns, mean of three experiments, Bars, S.E.M. **D.** Patient-derived (PD) mtNPM1 and MLL1-r AML cells were treated with the indicated concentrations of FHD-286 for 72 hours. Following this, cells were stained with TO-PRO-3 iodide and the % non-viable cells were determined by flow cytometry. **E-H.** MV4-11, MV4-11-MITR, OCI-AML3 and OCI-AML3-MITR cells were treated with the indicated concentrations of FHD-286 for 7 days. Following this, cells were assessed for the % expression and the mean fluorescent intensity (MFI) of CD11b by flow cytometry and morphologic features of differentiation. Columns, mean of three experiments, Bars, S.E.M.

### Effects of FHD-286 on chromatin occupancy of BRG1, enhancers and core regulatory circuitry in AML cells with MLL1r

To probe the basis of FHD-induced differentiation and lethal effects, we first determined the effects of FHD-286 on BRG1 occupancy, active enhancers and on core regulatory circuitry (CRC) in MOLM13 cells (16). ChIP-Seq analysis with anti-BRG1 antibody demonstrated that treatment with FHD-286 markedly reduced genome-wide occupancy of BRG1 on the chromatin (**Fig. 2A**). This was associated with decline in DNA binding sites of the key transcriptional regulators, including IRF8, PU.1 (SPI1) and its associated factor SPIB, and ETS1, at the loci with reduced BRG1 occupancy (**Fig. 2B**). FHD-286 caused a significant (p < 0.05) log2 fold-reduction in BRG1 occupancy at the loci active in LSCs and in signaling for cell growth, including WNT2B/5B, TCF4, LRP5, CLEC12A, CD244, GFIB, CSF, CSF1R, DUSP5 and CCND2 (**Fig. 2C**). ChIP-Seq with anti-H3K27Ac antibody that marks transcriptionally active enhancers and promoters showed genome-wide reduction in H3K27Ac occupancy (**Fig. S2A**). Specifically, the ROSE analysis showed FHD-286 treatment caused a loss of H3K27Ac peaks at the super-enhancers for ETS2, DNMT3B and RARA, and reduced H3K27Ac peak density at JUND, IRF8, JMJD1C and SPIB (**Figs. 2D** and **2E**). This was associated with a reduction in the CRC score from 5.81 to 4.71 and reduced numbers of participatory master regulators from 47 to 31, with the loss of activity of the master regulators KLF4, EGR3, WT1 and PRDM1 (**Fig S2B**) (16). In contrast, the ranking for the super-enhancers of GFI1, SPI1 (PU.1), CEBPA, IFNGR1, CD68 and HEXIM1 increased (**Figs. 2D** and **2E**). Concomitantly, FHD-286 treatment also caused significant log2 fold-increase in H3K27Ac peaks at these loci as well as those of BCL2A1, CD14, FBOX32, CD86, CDKN1B, HMOX1, and HSP1A/B (p < 0.05) (**Fig. 2F**).

**Figure 2.**
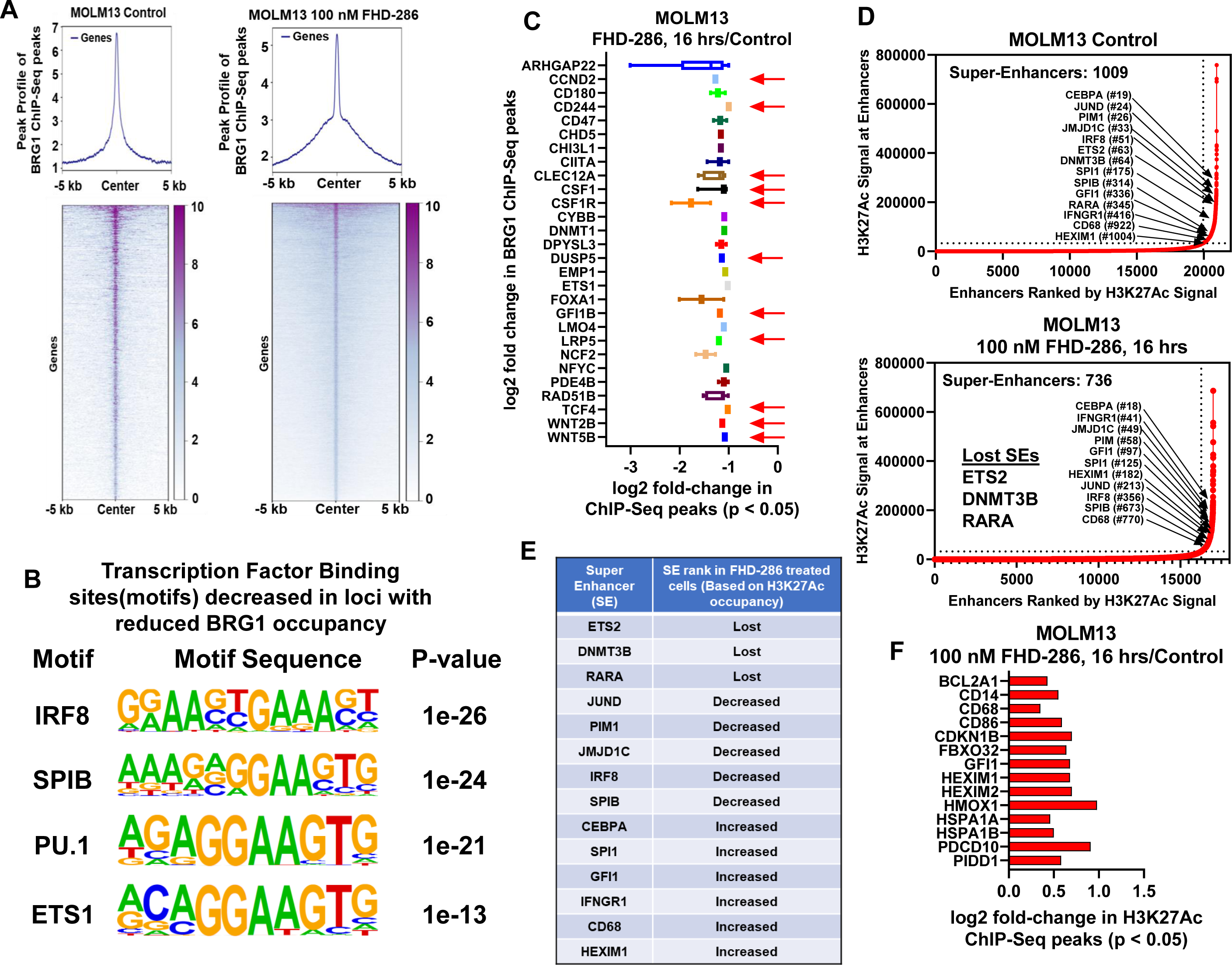
Treatment with FHD-286 depletes BRG1 occupancy on chromatin while increasing H3K27Ac occupancy on loci involved in differentiation and loss of cell viability. **A.** MOLM13 cells were treated with 100 nM of FHD-286 for 16 hours. ChIP-Seq analysis was conducted with anti-BRG1 antibody. Panel shows the genome wide peak profile and heat map of BRG1 binding at peak center +/- 5 kb resolution. **B**. Transcription Factor binding motifs decreased in loci with reduced BRG1 occupancy. The motif name, canonical binding motif and the p-value are shown. **C**. Log2 fold-decline of BRG1 binding at selected AML relevant loci in MOLM13 treated with 100 nM of FHD-286 for 16 hours. **D-E**. MOLM13 cells were treated with 100 nM of FHD-286 for 16 hours. ChIP-Seq analysis was conducted with anti-H3K27Ac antibody and ranked ordering of super enhancer (ROSE) analysis was performed. **F**. MOLM13 cells were treated with 100 nM of FHD-286 for 16 hours. ChIP-Seq analysis was conducted with anti-H3K27Ac antibody. Panel shows the log2 fold-increase in H3K27Ac occupancy on loci involved in differentiation and loss of viability in MOLM13 cells.

### Effects of FHD-286 on chromatin accessibility and transcriptome in AML cells

We next determined the genome-wide effects of FHD-286 on chromatin accessibility and mRNA expressions in MOLM13 cells. Following treatment with FHD-286, genome-wide, concordant, log2 fold-perturbations (upregulated: 1887 genes; downregulated: 2289 genes) in bulk ATAC-Seq and RNA-Seq peaks are shown in **Fig. 3A**, with peak reductions at several loci, including those of CDK4/6, LAMP5, CD244, MEF2D, PRDM1, FOXA1, IL7R and CSF1R, as well as peak increases at the loci including those of TNFRSF4, XBP1, GATA2, HEXIM1 and FBXO32 (**Fig. 3B**). Gene set enrichment analysis (GSEA) based on the RNA-Seq peak density perturbations revealed that FHD-286 treatment caused negative enrichment of HALLMARK gene-sets of MYC targets, MTORC1 signaling, inflammatory-response, IL6/JAK/STAT3 signaling, IFNγ-response, oxidative phosphorylation, and unfolded protein response, whereas gene-sets of TGFβ signaling and TNFα signaling via NFκB were positively enriched (**Figs. 3C** and **3D**). Notably, FHD-286 treatment caused log2 fold-decline of MYC, SPI1, IRF8, IL7R, MEF2D, CD180, CD44 and CSFR1 but increase in mRNA expressions of HEXIM1, PUMA, HMOX1, GATA2, XBP1 and TNFRSF4 (**Fig. 3E**). Following FHD-286 treatment, **Figs. S3A** to **S3C** show the heat map of RNA-Seq determined, ≥1.25-fold perturbations (p < 0.05) in mRNA expressions, negatively enriched mRNA gene-sets of the REACTOME pathways, as well as downregulation of specific MYC targets (e.g., SLC19A1) (32). FHD-286-mediated downregulation of specific mRNA transcript isoforms of MYC, PU.1 and BCL2 are shown in **Figs. S3D** to **S3F**. Treatment with FHD-286 also perturbed the expressions of ERVs (endogenous retroviruses). **Figs. S3G** and **S3H** show the heat map of ERV transcripts upregulated or downregulated and the log2 fold-change in specific ERVs impacted, following FHD-286 treatment. Utilizing AML bone marrow aspirate (BMA) cells of a patient with AML expressing mtNPM1 and FLT3-ITD, we also conducted single cell (sc) multiomics analyses involving ATAC-Seq and RNA-Seq in untreated and FHD-286-treated cells. The UMAP (Uniform Manifold Approximation and Projection plot) revealed that FHD-286 treatment reduced cell events in the MEP (megakaryocyte-erythroid progenitor) cluster, denoting reduction in the MEP population of cells, whereas the cell numbers of macrophages/monocytes increased (**Fig. 4A and 4B**). Exposure to FHD-286 caused significant log2 fold-decline in ATAC-Seq peaks at the MYC, CSF1R, SPI1, MEF2C, HOXA9 and RUNX1 loci, including reduced binding motifs for the TFs ETS1, ERG, PU.1, SPIB and RUNX1, but increase in ATAC-Seq peaks at CBX1/5, KMD6B/JMJD3 and BBC3 (**Figs. 4C** **and S4A**). Notably, ATAC-Seq peak density significantly decreased at the SPI1 locus as shown in the Integrated Genome Viewer (IGV) plot in **Fig. S4B**. Following FHD-286 treatment, scRNA-Seq revealed negative enrichment of mRNAs of gene-sets belonging to MYC targets, inflammatory response, IL6/JAK/STAT3 signaling, E2F targets, G2/M checkpoint and DNA repair, like what was observed in FHD-286-treated MOLM13 cells (**Fig. 4D**). The volcano plot of FHD-286-mediated mRNA perturbations is shown in **Fig. 4E**. Similar findings were observed in a PD AML sample with MLL-AF9 expression treated with FHD-286 for 16 hours (**Figs S4C to S4E**).

**Figure 3.**
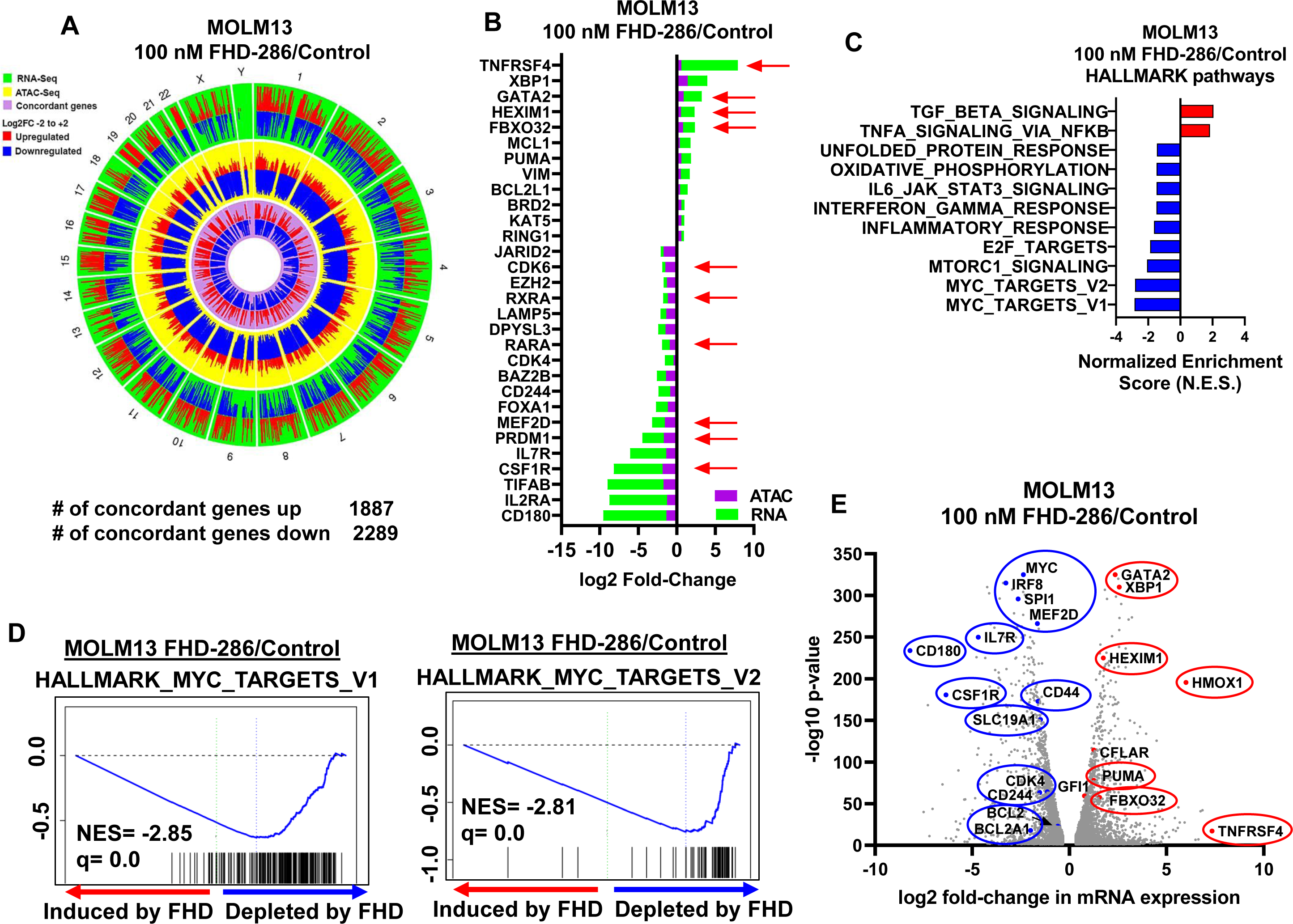
Treatment with FHD-286 concordantly alters chromatin accessibility and mRNA expression in MLL1-rearranged AML cells with reduction in the enrichment scores for MYC, mTORC1, E2F, Interferon-gamma, IL6-JAK-STAT3, as well as of inflammatory response and oxidative phosphorylation gene sets. **A-B.** MOLM13 cells were treated with 100 nM of FHD-286 for 16 hours as biologic replicates. Bulk nuclei were isolated for ATAC-Seq analysis and total RNA was isolated and utilized for RNA-Seq analysis. RNAs and diffReps-determined ATAC-Seq peaks with ≥ 1.25 fold-change and p <0.05 were utilized for the concordance analysis. Circos plot (**A**) and log2 fold-changes (**B**) of selected, concordant ATAC-Seq and mRNA expression alterations in FHD-286-treated MOLM13 cells. **C**. Gene set enrichment analysis of FHD-286-treated MOLM13 cells compared to HALLMARK pathways. Normalized enrichment scores are shown. All q-values are < 0.1. **D**. Enrichment plot of FHD-286-treated MOLM13 cells compared to HALLMARK_MYC_TARGETS_V1 and HALLMARK_MYC_TARGETS_V2. **E**. Volcano plot (log2 fold-change versus –log10 p-value) of RNA-Seq determined mRNA expression changes in FHD-286-treated MOLM13 cells.

**Figure 4.**
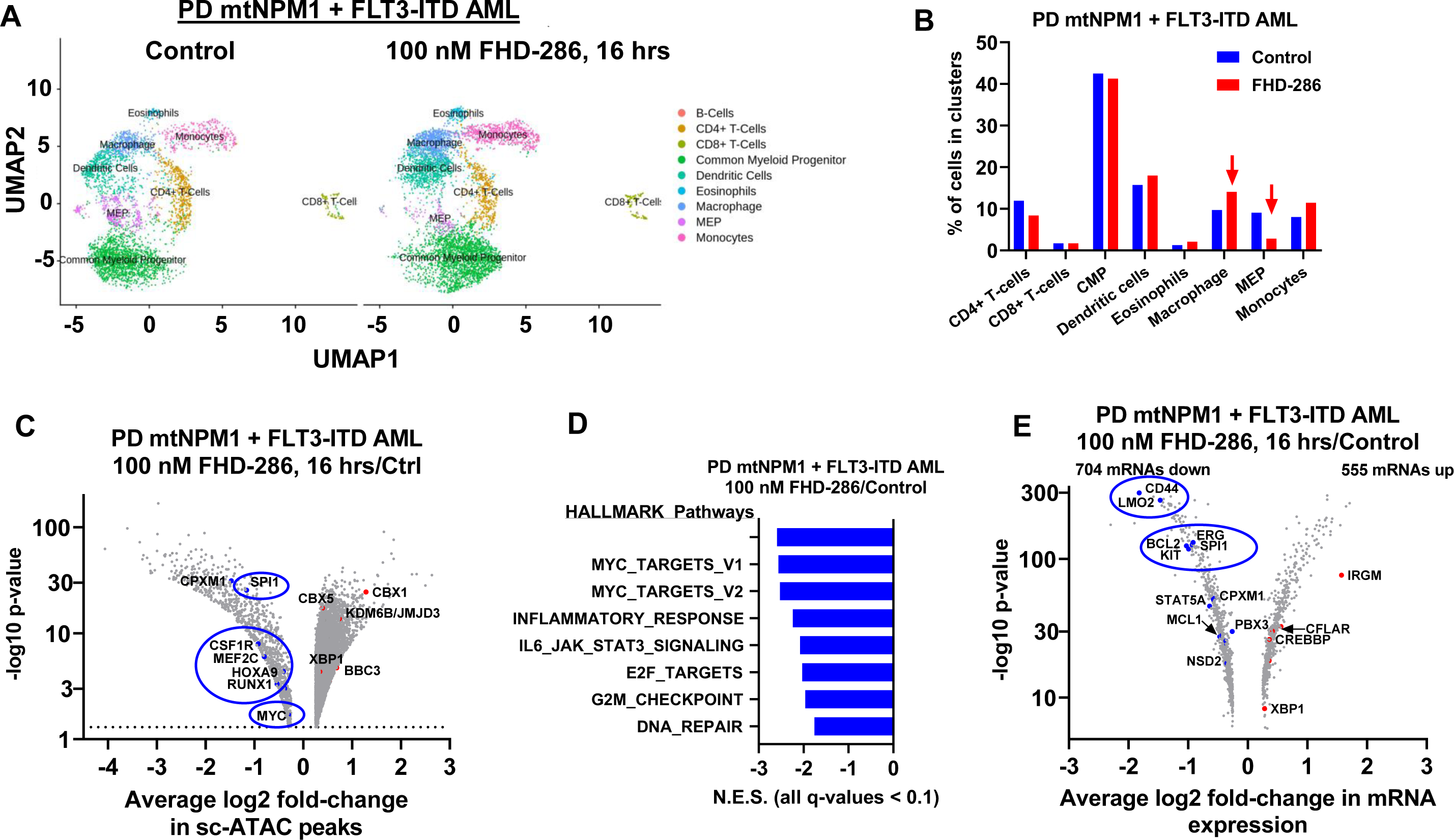
Treatment with FHD-286 depletes MEP cells and reduces chromatin accessibility and enrichment scores of MYC TARGET genes in the CMP cluster of patient-derived mtNPM1 + FLT3-ITD expressing AML cells. **A.** Patient-derived mtNPM1-expressing AML cells from a bone marrow aspirate were treated with 100 nM of FHD-286 for 16 hours. Multiomics (combined sc-ATAC-Seq and sc-RNA-Seq) analyses were performed on isolated nuclei. The UMAP plot shows the SingleR-determined composition of the individual cell clusters in the control and FHD-286-treated AML cells. **B**. Percentage of each cell type in the clusters of cells from control and FHD-286 treated cells. Arrows indicate clusters with increased or decreased numbers of cells in the FHD-286-treated sample compared to the control sample. **C**. Volcano plot of sc-ATAC-Seq peaks in the CMP cluster with ≥1.25 fold-change up or down and p< 0.05 following treatment with FHD-286. **D**. Gene set enrichment analysis of FHD-286-treated cells over control cells. All q-values are less than 0.1. **E**. Volcano plot of sc-RNA-Seq expression changes (≥1.25 fold-change and p< 0.05) in the CMP cluster following treatment with 100 nM FHD-286 for 16 hours compared to control cells.

### FHD-286-mediated protein expression alterations include c-Myc, PU.1 and CDK6 in phenotypically defined AML cells with MLL1r or mtNPM1

To determine FHD-286 induced protein expression perturbations signature, we next conducted mass spectrometry on cell lysates from untreated or FHD-286-treated MOLM13 and PD AML cells. **Fig. S5A** lists the overlapping 75 proteins that showed a decline and the 52 proteins that were upregulated following FHD-286 treatment, with the log2 fold-change in each protein level. Out of these protein expression perturbations, a signature of selected, overlapping, depleted (15 proteins) and increased (7 proteins) expression alterations (≥ 1.25-fold and p < 0.05) due to FHD-286 treatment are depicted in **Fig. 5A**. These proteins are involved in AML cell growth, cell cycle progression and cell death. Altered protein expressions included reduced levels of c-Myc, PU.1, CD44, SLC19A1, MATK and CSF2RA, as well as increased levels of CDKN1B, HMOX1, DNASE2 and NEU1. We next compared the protein signature of the overlapping 75 depleted and 52 induced proteins in both AML samples against REACTOME pathways. Following FHD-286 treatment, the sets of proteins belonging to the ‘reactomes’ of RNA pol II transcription and signal transduction were reduced, whereas the proteins belonging to the ‘reactome’ of immune stimulation showed increased expressions (**Figs. 5B** **and S5B**). In MOLM13 cells, the volcano plot of significant, log2 fold-increased or decreased proteins are shown in **Fig. S5C,** whereas **Fig. S5D** presents the specific downregulation of proteins within the HALLMARK gene-set of MYC targets but upregulation of proteins within the gene-set of inflammatory response. **Fig. S5E** demonstrates the volcano plot of significantly up- or down-regulated proteins in the FHD-286-treated versus untreated PD AML cells with mtNPM1 and FLT3-ITD. It depicts log2 fold-decline in the protein expressions of SPI1, c-Myc, CD180, CDK4, CD44 and SLC19A1 with concomitant increase in AIF1, IRF8, CASP9, HMOX1 and p27 levels. Additionally, gene set enrichment analysis of the protein expression signature demonstrated the negative enrichment of gene-sets belonging to HALLMARK MYC Targets, E2F targets and G2M checkpoint **(Fig. S5F**). We also confirmed by immunoblot analyses FHD-286-mediated protein expression changes in MOLM13 cells. It showed that, in the FHD-286-treated versus untreated MOLM13 cells, there was a time-dependent increase in TP53, p21, p27, PUMA and CD11b, but decline in the protein levels of c-Myc, BCL2, and FLT3 (**Fig. S5G**). Utilizing cocktails of rare-metal element-tagged antibodies, we next performed CyTOF analyses on two samples each of PD AML cells with MLL1r (Sample #24 and Sample #18 in the oncoplot) or mtNPM1 (Sample #15 and Sample #19 in the oncoplot). **Fig. 5C** shows that, in the phenotypically defined AML stem cells (high expression of CLEC12, CD123, CD99, and CD33 but low expression of CD11b), compared to the untreated controls, FHD-286 treatment depleted the protein expressions of BRG1, PU.1, RUNX1, c-Myc, CDK6 and MEF2C, but increased expressions of cleaved PARP and p-H2AX. Following FHD-286 treatment, there was also a significant decline in the phenotypically defined stem cell frequency in the AML samples with MLL1r but not in the sample with mtNPM1 (**Fig. 5D**).

**Figure 5.**
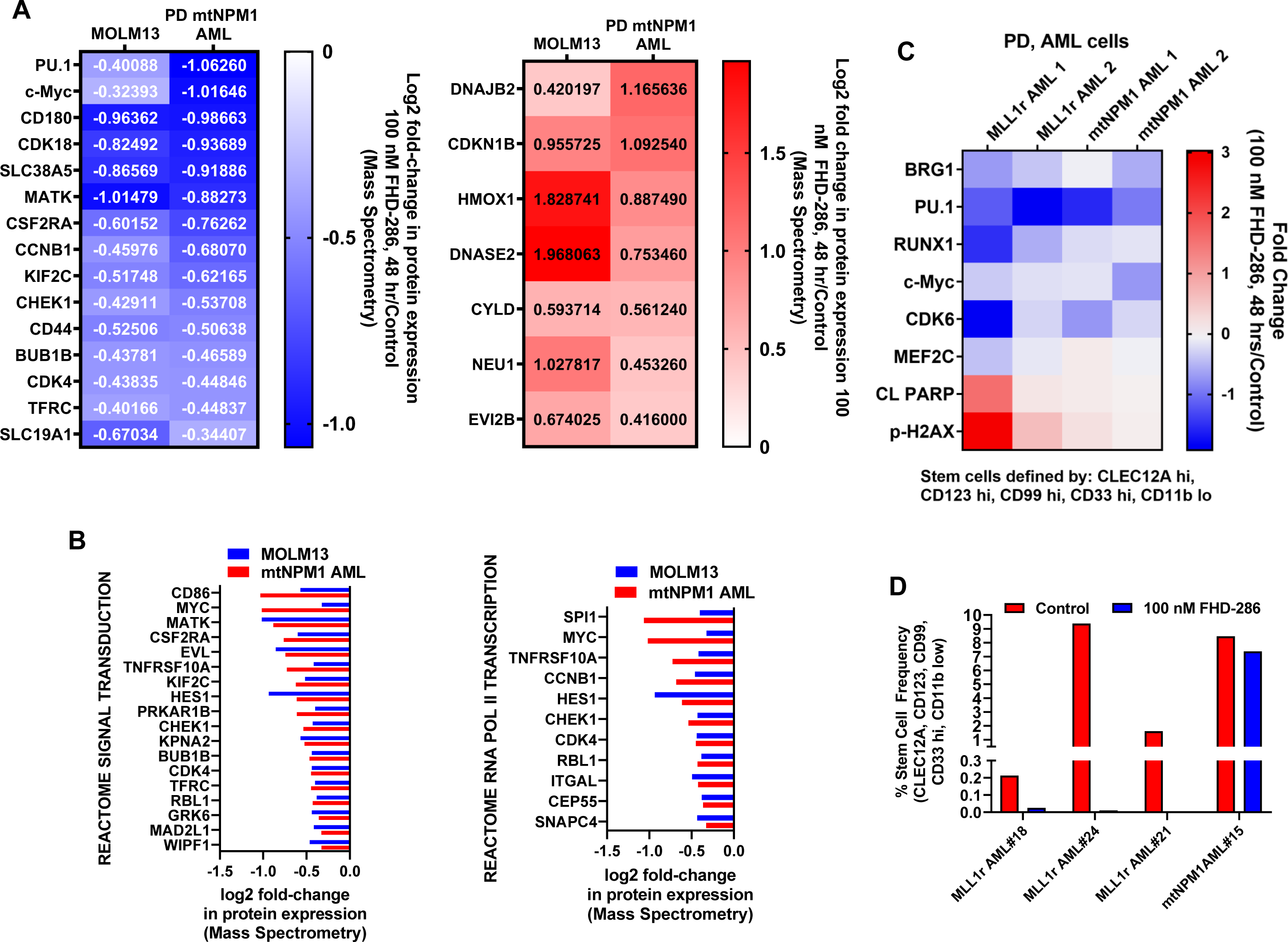
Treatment with FHD-286 significantly depleted c-Myc and PU.1 expression in bulk AML cells and phenotypically defined AML stem cells with MLL1r or mtNPM1. **A**. MOLM13 and PD mtNPM1 expressing AML cells were treated with 100 nM of FHD-286 for 48 hours. Total proteome profiling was conducted by mass spectrometry analysis. The heat map shows selected overlapping depleted and induced protein expressions with a fold change greater than 1.25 and a p-value <0.05. **B**. Log2 fold-decline in protein expressions in FHD-286-treated MOLM13 and PD mtNPM1-expressing AML cells compared to REACTOME_SIGNAL_TRANSDUCTION and REACTOME_RNA_POLII_TRANSCRIPTION pathways. **C**. Patient-derived MLL1r and mtNPM1 AML cells were treated with 100 nM of FHD-286 for 48 hours. CyTOF analyses were conducted utilizing cocktails of rare metal element tagged antibodies. The heat map shows the fold change (FHD-286 treated over control) of depleted and induced proteins in phenotypically defined AML stem cells (CLEC12A hi, CD123 hi, CD99 hi, CD33 hi and CD11b low). **D**. Percent stem cell frequency of control and FHD-286 treated patient-derived MLL1r and mtNPM1 expressing AML cells.

### In vivo efficacy of FHD-286 *to* target AML-initiating stem cells

We next determined the in vivo efficacy of FHD-286 against AML-initiating LSCs, utilizing a PDX model of luciferase-transduced AML with mtNPM1 and FLT3-ITD/TKD. Following ex vivo treatment with 10 or 30 nM of FHD-286 for 96 hours, the PDX cells were tail-vein infused and engrafted in NSG mice. After one, two or 4 weeks, AML burden, represented by the total bioluminescence flux, was determined. **Figs. 6A, 6B** **and S6A** demonstrate that, compared to the untreated control, the ex vivo exposure to FHD-286 led to a significant in vivo decline in the AML burden and improvement in the median and overall survival of the NSG mice. In a separate study, following infusion and engraftment of the AML PDX cells, mice were treated for 5 weeks with FHD-286 or vehicle control. As shown in **Figs. 6C to 6E**, compared to the vehicle treated mice, treatment with FHD-286 caused significant decline in the AML burden and spleen size in the NSG mice. FHD-286 treatment for 5 weeks also led to a significant improvement in the survival of NSG mice (**Fig. S6B**). In an aggressive PDX model of AML with mtNPM1 and FLT3-ITD/FLT3-F691L, 5-weeks of treatment with FHD-286 also yielded a significant improvement in the survival of the mice (**Fig. S6C**). After 5 weeks of FHD-286 treatment of the engrafted PDX of AML with mtNPM1 and FLT3-ITD/TKD, BM or spleen cells were harvested from the cohorts of mice treated with the vehicle control or FHD-286, and the cells were re-infused separately into cohorts of mice which did not receive any further treatment with FHD-286. **Figs. 6F and 6G** demonstrate that mice re-implanted with the AML cells from the previously FHD-286-treated mice exhibited significantly lower AML burden accompanied by improved survival, as compared to those re-implanted with cells from the previously vehicle-treated cohort (**Fig. 6H**). Taken together, these findings reveal that FHD-286 treatment significantly attenuates AML-initiating stem-progenitor cells in the AML PDX models. We also determined the host toxicity of FHD-286 in the immune-competent C57/BL6 mice. Following two weeks of treatment with 1.5 mg/kg of FHD286, there was no significant decline in the mouse wight, white cell counts or hematocrit (**Figs. S6D and S6E**). However, FHD treatment reduced the platelet counts, which recovered upon withdrawal of the drug during the 2-weeks of the recovery period (**Fig. S6E**).

**Figure 6.**
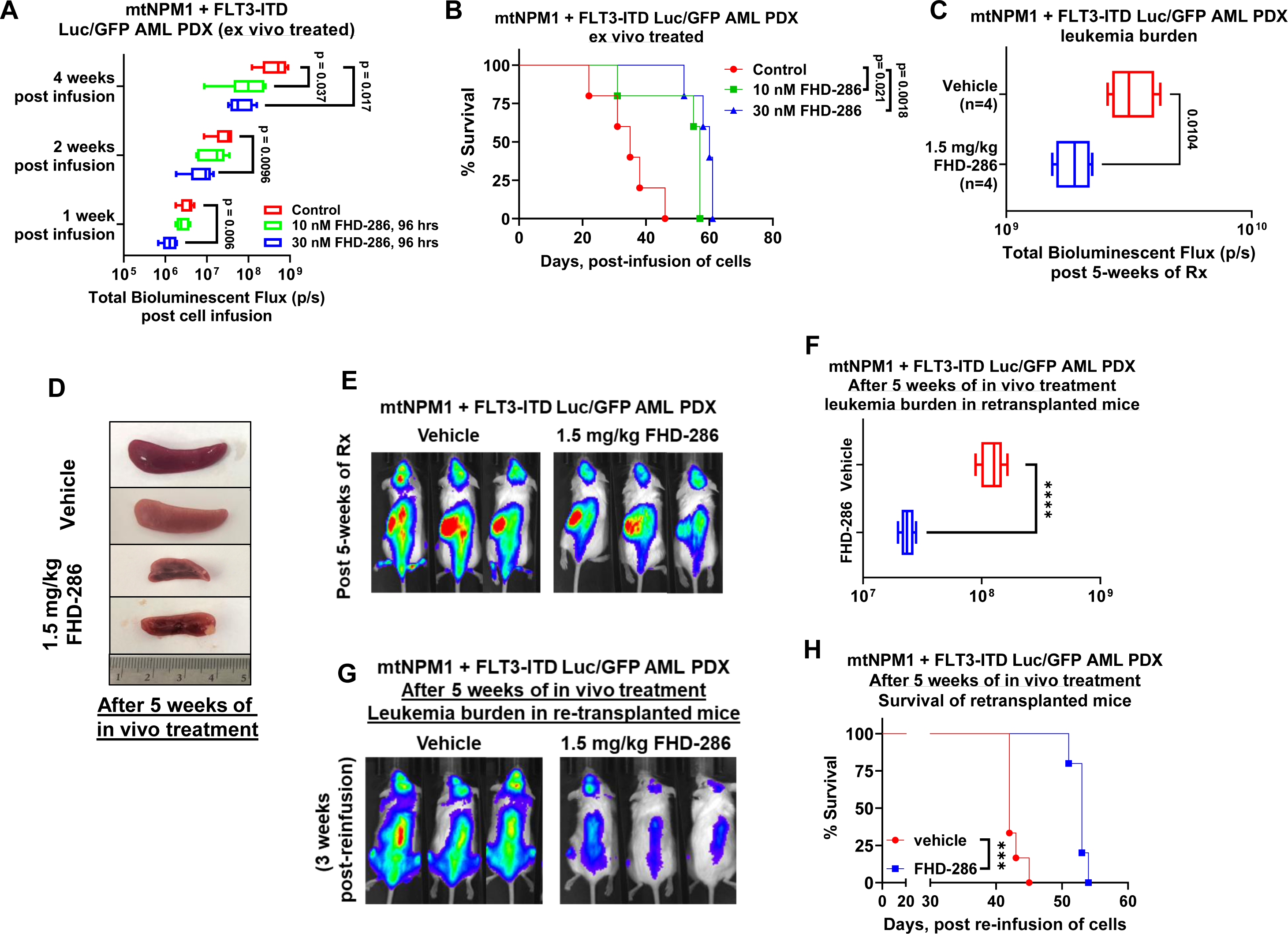
Treatment with FHD-286 exhibits in vivo efficacy against AML initiating stem cells. **A**. Patient-derived mtNPM1 + FLT3-ITD Luc/GFP cells (AML#1 in the oncoplot) were ex vivo treated with 10 and 30 nM of FHD-286 for 96 hours. Following this, equal numbers of cells (2.5e6 cells/mouse) were tail vein infused into pre-irradiated (2.5 Gy) NSG mice (n=5 per cohort). Mice were monitored daily for symptoms of acute leukemia. Luciferase signal in the mice was determined by IVIS (Xenogen) imaging at one, two and four weeks post-infusion of AML cells. The box plots show the total bioluminescent flux (photons/second) at one, two, and four weeks post infusion of the AML cells. Significance between cohorts was determined by a two-tailed, unpaired t-test utilizing GraphPad Prism V9. P-values less than 0.05 were considered significant. **B**. Kaplan-Meier survival curve of NSG mice infused with ex vivo treated patient-derived mtNPM1 + FLT3-ITD Luc/GFP cells. Significance between cohorts was determined by a Mantel-Cox log-rank test. P-values less than 0.05 were considered significant. **C**. Patient-derived mtNPM1 + FLT3-ITD Luc/GFP cells (2.5e6 cells/mouse) were tail vein infused into pre-irradiated (2.5 Gy) NSG mice (n=4 per cohort). Mice were monitored for 5 days and leukemia engraftment was documented by IVIS imaging. Mice were randomized to equivalent bioluminescence and treated with vehicle or 1.5 mg/kg of FHD-286 for 5 weeks. The box plots show the total bioluminescent flux (photons/second) at five weeks post infusion of the AML cells. Significance between cohorts was determined by a two-tailed, unpaired t-test utilizing GraphPad Prism V9. P-values less than 0.05 were considered significant. **D**. Post 5 weeks of treatment when vehicle mice required euthanasia, all mice were sacrificed and the spleens and bone marrow were harvested. The panel shows two representative spleens from vehicle and 1.5 mg/kg FHD-286 treated mice. **E**. Viable human AML cells from the spleens and bone marrow of vehicle and FHD-286-treated mice were re-infused into pre-irradiated (2.5 Gy) NSG mice (n=6 per cohort). The box plots show the total bioluminescent flux (photons/second) three weeks post re-infusion of the AML cells. Significance between cohorts was determined by a two-tailed, unpaired t-test utilizing GraphPad Prism V9. P-values less than 0.05 were considered significant. **F**. Representative bioluminescent images of mice from panel C and E. **G**. Kaplan-Meier survival curve of NSG mice infused with equal numbers of previously in vivo treated patient-derived mtNPM1 + FLT3-ITD Luc/GFP cells. Significance between cohorts was determined by a Mantel-Cox log-rank test. P-values less than 0.05 were considered significant.

### Superior efficacy of combined therapy with FHD-286 with decitabine, venetoclax, BET inhibitor or Menin inhibitor in PDX models of AML with MLL1r or mtNPM1

We next determined the in vitro lethal activity of co-treatment with FHD-286 and decitabine, venetoclax, BETi and MI against AML cells with MLL1r or mtNPM1. Combined treatment with FHD-286 and decitabine induced synergistic lethality in MV4-11, MOLM13 and OCI-AML3 cells, with delta synergy scores of over 10 by the ZIP method (**Fig. S7A**). Additionally, co-treatment with FHD-286 and venetoclax, OTX015 or SNDX-50469 induced synergistic loss of viability in the AML cell lines and PD AML cells with MLL1r or mtNPM1, with mean delta synergy scores over 5 by the ZIP method (**Fig. 7A**). Notably, co-treatment with FHD-286 and OTX015 or SNDX-50469 also induced synergistic loss of viability in MV4-11-MITR and OCI-AML3-MITR cells that lacked Menin mutations (as above) (**Fig. 7A**). Moreover, treatment with FHD-286 alone or in combination with OTX015 or SNDX-50469 did not induce loss of viability in normal CD34 progenitor cells (less than 15% of control), indicating relative in vitro sparing of normal hematopoietic progenitors (**Figs. S7B and S7C**). Next, we evaluated the pre-clinical in vivo efficacy of FHD-286-based combinations in the PDX models of AML with MLL1r or mtNPM1. Following tail vein infusion and engraftment of luciferase-transduced PD AML cells expressing MLL-AF9 and mtFLT3, cohorts of NSG mice were treated with vehicle control or previously-determined safe doses of the drugs, i.e., FHD-286 and/or venetoclax or decitabine or the BET inhibitor OTX015. In the MLL-AF9 and mtFLT3 harboring AML PDX cells the co-mutations and their variant allelic frequencies detected by NextGen sequencing are listed in **Table S1**. Notably, **Fig. 7B to 7E** demonstrates that, compared to treatment with vehicle control, monotherapy with FHD-286, decitabine or OTX015, but not venetoclax, significantly reduced the AML burden as well as improved the survival of the NSG mice. Importantly, as compared to treatment with each drug alone, co-treatment with FHD-286 and decitabine or venetoclax or OTX015 was significantly more effective in reducing the AML burden and improving the survival of NSG mice (**Figs. 7B to 7E** **and S7D**). We also determined the in vivo efficacy of co-treatment with FHD-286 and BETi or MI in a PDX model of AML cells with mtNPM1 and FLT3-ITD. The co-mutations and their variant allelic frequencies detected in this AML PDX cells are also shown in **Table S1**. Here, combined therapy with FHD-286 and the safe doses of OTX015 or SNDX-5613 reduced significantly more AML burden and yielded significantly better survival than treatment with each drug alone or vehicle control (**Figs. 7F to 7I)**. These findings highlight the superior in vivo efficacy of FHD-286 monotherapy as well as the combination therapy with not only the established anti-AML agents such as decitabine or venetoclax but also with MI or BETi against the PDX models of MLL1r or mtNPM1-expressing AML cells.

**Figure 7.**
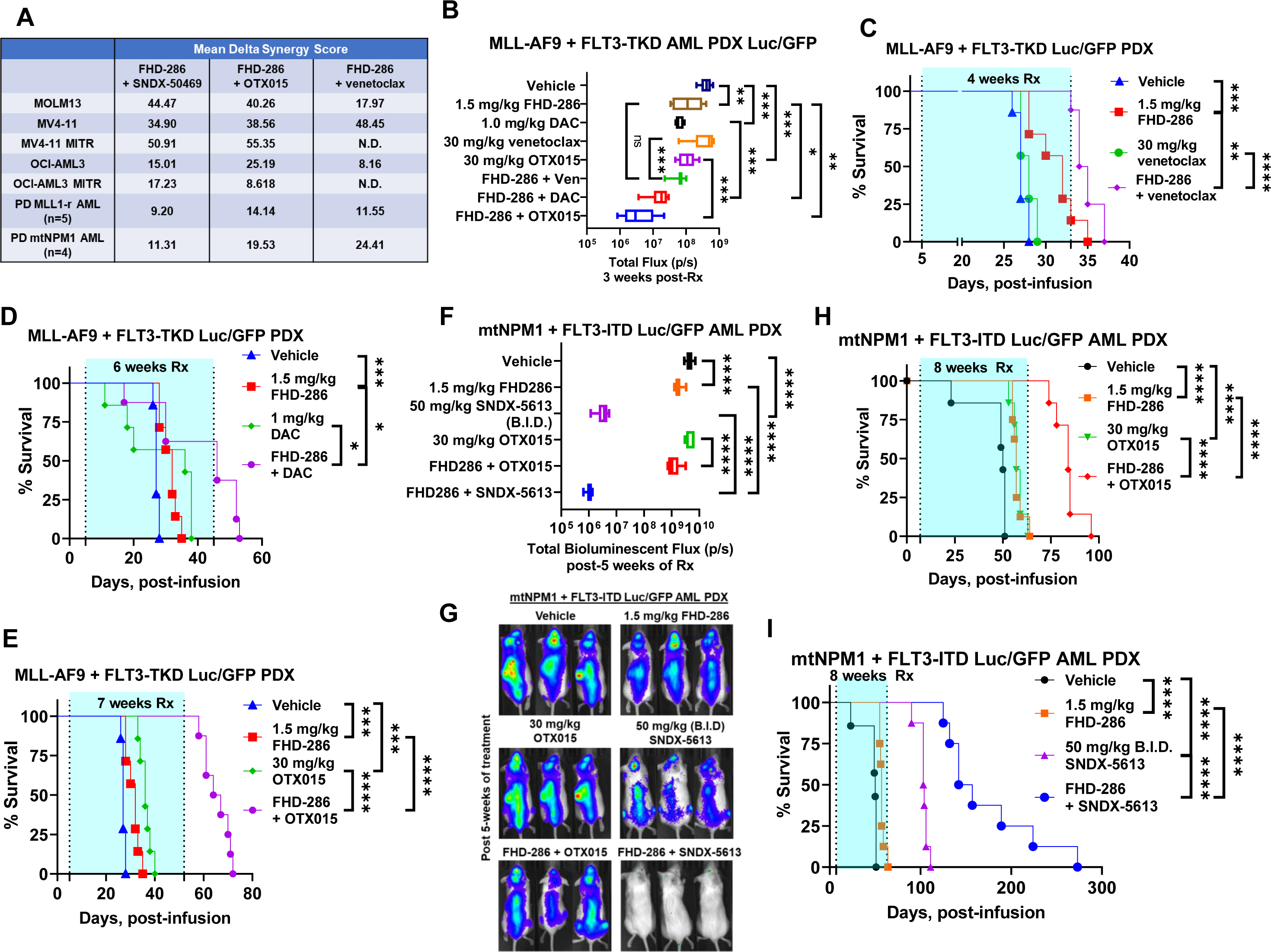
Treatment with FHD-286-based combinations exerted synergistic *in vitro* lethality in cultured and patient-derived AML cells expressing MLL-r or mtNPM1 with or without mtFLT3 and reduced leukemia burden and significantly improved survival of NSG mice bearing MLL-r or mtNPM1-expressing AML xenografts. **A.** MOLM13, MV4-11, MV4-11-MITR, OCI-AML3, OCI-AML3-MITR and patient-derived MLL1r or mtNPM1-expressing AML cells were treated with FHD-286 (dose range: 10 nM to 250 nM) and Menin inhibitor SNDX-50469 (dose range: 50 nM to 1000 nM), BET inhibitor OTX015 (dose range: 50 nM to 250 nM) or venetoclax (dose range: 10 nM to 100 nM) for 72-96 hours. At the end of treatment, the % non-viable cells was determined by staining with To-Pro-3 iodide and flow cytometry analysis. Delta synergy scores were determined by the ZIP method within the SynergyFinder web application. Synergy scores >1.0 indicate a synergistic interaction of the two agents in the combination. Panel shows the mean Delta Synergy score for each FHD-286 based combination in the cell lines and PD AML cells. **B**. Total photon counts [flux] (determined by bioluminescent imaging) in NSG mice engrafted with luciferized MLL-AF9 + FLT3-TKD AML PDX cells (AML#6 in the oncoplot) and treated for 3 weeks with FHD-286 and/or venetoclax, decitabine or OTX015 at the indicated doses. **C**. Kaplan-Meier survival plot of NSG mice engrafted with luciferized MLL-AF9 + FLT3-TKD AML PDX cells and treated with 1.5 mg/kg of FHD-286 (daily x 5 days, P.O.) and/or 30 mg/kg of venetoclax (daily x 5 days, P.O.) for 4 weeks. Significance was calculated by a Mantel-Cox log-rank test. **D**. Kaplan-Meier survival plot of NSG mice engrafted with luciferized MLL-AF9 + FLT3-TKD AML PDX cells and treated with 1.5 mg/kg of FHD-286 (daily x 5 days, P.O.) and/or 1 mg/kg of DAC (Days 1-5 days only, IP) for 6 weeks. Significance was calculated by a Mantel-Cox log-rank test. **E**. Kaplan-Meier survival plot of NSG mice engrafted with luciferized MLL-AF9 + FLT3-TKD AML PDX cells and treated with 1.5 mg/kg of FHD-286 (daily x 5 days, P.O.) and/or 30 mg/kg of OTX015 (daily x 5 days, P.O.) for 7 weeks. Significance was calculated by a Mantel-Cox log-rank test. **F**. Total photon counts [flux] (determined by bioluminescent imaging) in NSG mice engrafted with luciferized mtNPM1 + FLT3-ITD PDX cells and treated for 3 weeks with FHD-286 and/or SNDX-5613 or OTX015 at the indicated doses. **G**. Representative bioluminescent images of mice from panel F. **H-I**. Kaplan-Meier survival plot of NSG mice engrafted with luciferized mtNPM1 + FLT3-ITD PDX cells and treated with 1.5 mg/kg of FHD-286 (daily x 5 days, P.O.) and/or 30 mg/kg of OTX015 (daily x 5 days, P.O.) or SNDX-5613 (50 mg/kg, B.I.D. x 5 days, P.O) for 8 weeks. Significance between cohorts was determined by a Mantel-Cox log-rank test.

## Discussion

The results presented in this manuscript provide compelling evidence regarding the dependency of AML cells with MLL rearrangements (MLL1r) or mutant NPM1 (mtNPM1) on BRG1/BRM for the differentiation arrest, growth and survival, as well as on their sensitivity to FHD-286-mediated inhibition of BRG1/BRM in inducing differentiation and loss of viability. These findings have significant implications for understanding the molecular mechanisms underlying AML pathogenesis and leukemia initiation potential of AML stem-progenitor cells. They also highlight the potential FHD-286-based combination therapeutic strategies involving MI, BETi and the standard anti-AML agents, such as decitabine and venetoclax, against AML with MLL1r or mtNPM1.

Utilizing DepMap findings, we determined here that SMARCA4 (BRG1) is a dependency in the various AML cell lines, including those expressing MLL fusion proteins (MLL-FP) and mutant NPM1 (mtNPM1). This underscores the potential therapeutic significance of targeting SMARCA4 in the specific AML subtypes represented by the cell lines. This dependency on SMARCA4 for survival also suggests its central role in the maintenance of AML cells and a promising therapeutic target (2, 4, 6, 7). Our findings highlight FHD-286, a clinical-grade dual BRG1 and BRM inhibitor, as a potential effective therapeutic agent for AML with MLL1r or mtNPM1. FHD-286 treatment induces the expression of the CD11b marker and morphological features of differentiation, while also significantly reducing AML cell viability, including of AML cells with TP53 mutations. Importantly, FHD-286 appears to be selectively more toxic against the AML cells, sparing normal CD34+ progenitor cells. Previous reports have highlighted that targeting BRM is synthetic-lethal in BRG1 deficient cancers (8, 33). This unique dependency on a subunit paralog or the downstream targets or pathways should be targeted by FHD-286, since it simultaneously inhibits the catalytic activity of BRG1 and BRM, which would also potentially inhibit all three BAF complexes in AML stem-progenitor cells (6,34). This may also explain our findings that, regardless of the BRG1 levels, FHD-286 exerted superior in vitro and in vivo efficacy against the AML stem-progenitor cells. Specific targeting of BRG1/BRM with a protein degrader is also likely to yield similar outcome and needs to be evaluated in this setting (35).

Findings of ChIP-Seq analyses revealed that FHD-286 treatment leads to a reduction in BRG1 and H3K27Ac occupancy across the genome, impacting the binding sites of key transcriptional regulators such as IRF8, PU.1, SPIB, and ETS1. These changes disrupted the core regulatory circuitry and affected gene-expressions associated with AML cell growth, maintenance and survival. Specifically, FHD-286 treatment had a profound impact on the transcriptome and proteome of AML cells, leading to widespread changes in gene expressions, affecting pathways including those related to MYC targets, inflammatory responses, cell cycle regulation (2, 7). These downstream effects may be directly related to reduced BRG1 binding to its chromatin targets and through indirect gene regulation beyond the effects on BRG1-binding to the chromatin (2, 3). Our studies involving mass spectrometry and immunoblot analyses also highlighted specific protein expression alterations, including the downregulation of c-Myc, c-Myb, PLK1, PU.1, FLT3 and CDK4, which are crucial factors blocking differentiation and promoting growth and survival of AML cells with MLL1r or mtRUNX1. Results of the CyTOF studies are especially important in demonstrating that FHD-286 treatment attenuates a signature of protein expressions, including those of BRG1, c-Myc, PU.1, CDK6 and RUNX1 in the phenotypically defined AML stem-progenitor cells expressing MLL1-FP or mtNPM1. Concomitantly, based on these data, FHD-286 treatment also reduced the AML stem-progenitor cell frequency. Additionally, in CyTOF studies, the notable down-regulation of BRG1 by FHD-286 treatment could be due to FHD-286-mediated inhibition followed by destabilization of the BRG1 protein. Taken together, these findings provide valuable insights into the molecular mechanisms through which FHD-286 overcomes differentiation block, undermines AML-initiating potential and loss of viability in AML stem-progenitor cells.

Our findings also address the issue of drug tolerance-resistance, a common challenge in AML treatment. Notably, FHD-286 remained effective in AML cells that had developed resistance to Menin inhibitors, suggesting that it could be valuable in overcoming drug tolerance/resistance through non-genetic or adaptive mechanisms (10, 31, 36). By inhibiting BAF complex mediated chromatin remodeling and thereby the enhancer activities of master regulators such as c-Myc and PU.1, treatment with FHD-286 may overcome the AML progenitor cell-plasticity and their escape through dedifferentiation or through a phenotypic switch (31, 37, 38). This finding opens new possibilities for including FHD-286 in the targeted combination therapies for combating drug-tolerant/resistant AML stem-progenitor cells. As shown in the PDX model of AML with mtNPM1 and FLT3-ITD/TKD, FHD-286 targeted AML-initiating stem-progenitor cells, as well as significantly reduced AML burden and improved overall survival in these AML models. Importantly, FHD-286 also showed a favorable safety profile in the host mice, further supporting its potential clinical utility. Combination therapies involving FHD-286 along with the established AML drugs, including decitabine, venetoclax, BET inhibitor, or Menin inhibitor, exhibited superior efficacy in reducing the AML burden and enhancing the survival in AML PDX models. These results suggest the potential for including FHD-286 in combination with standard and novel targeted therapies to improve the outcomes in AML with MLL1r or mtNPM1.

In conclusion, the findings presented here provide a comprehensive understanding of the role of SMARCA4 (BRG1) in AML cells with MLL1r or NPM1. The dual BRG1/BRM ATPase inhibitor FHD-286 emerges as a promising therapeutic agent against the AML stem-progenitor cells of these AML subtypes, inducing differentiation, overcoming drug resistance, and relatively sparing the normal hematopoietic progenitor cells. The insights by this study into epigenetic alterations, transcriptomic changes, and protein expression alterations shed light on the molecular mechanisms underlying the therapeutic efficacy of FHD-286. Additionally, the in vivo efficacy of FHD-286-based combinations highlighted here further supports the clinical development of FHD-286 as a novel therapeutic approach in the treatment of AML with MLL1r or mtNPM1 in the MRD or relapsed AML setting.

## Materials and Methods

### Reagents and antibodies

FHD-286 was obtained under a material transfer agreement with Foghorn Therapeutics (Cambridge, MA). Venetoclax, OTX015, decitabine, SNDX-50469 and SNDX-5613 for in vivo studies were obtained from MedChem Express (Monmouth Junction, NJ). All compounds for in vitro studies were prepared as 10 mM stocks in 100% DMSO and frozen at -80°C in 5-10 µL aliquots to allow for single use, thus avoiding multiple freeze-thaw cycles that could result in compound decomposition and loss of activity. Anti-c-Myc [RRID: AB_1903938], anti-PUMA [RRID:AB_2797920], and anti-p21 Waf1/Cip1 [RRID:AB_823586] antibodies were obtained from Cell Signaling Technologies (Beverly, MA). Anti-BRG1 [Cat# ab110641, RRID:AB_10861578] Anti-FLT3 [Cat# ab245116] and anti-CD11b [RRID:AB_2650514] antibodies were obtained from Abcam (Cambridge, MA). Anti-BCL2 [RRID: AB_626733], and anti-GAPDH [RRID: AB_627679] antibodies were obtained from Santa Cruz Biotechnologies (Dallas, TX). Anti-p27 [RRID:AB_397636] and anti-p53 [RRID:AB_395348] antibodies were obtained from BD Biosciences (Franklin Lakes, NJ). H3K27Ac [RRID:AB_2793305] antibody was obtained from Active Motif (Carlsbad, CA).

### Cell lines and cell culture

MOLM13 [DSMZ Cat# ACC-554, RRID: CVCL_2119] and OCI-AML3 [DSMZ Cat# ACC-582, RRID:CVCL_1844] cells were obtained from the DSMZ (Braunschweig, Germany). MV4-11 [ATCC Cat# CRL-9591, RRID:CVCL_0064] cells were obtained from the ATCC (Manassas, VA). MOLM13 cells with isogenic TP53 mutations [R175H, R248Q and TP53-KO] were a gift from Dr. Benjamin L. Ebert (Dana Farber Cancer Center, Boston, MA). HEK-293T [RRID:CVCL_0063] cells were obtained from the Characterized Cell Line Core Facility at M.D. Anderson Cancer Center, Houston TX. All experiments with cell lines were performed within 6 months after thawing or obtaining from ATCC or DSMZ. MOLM13 and OCI-AML3 cells were cultured in RPMI-1640 media with 20% FBS, 1% penicillin/streptomycin and 1% non-essential amino acids. MV4-11 cells were cultured in ATCC-formulated IMDM media with 20% FBS, 1% penicillin/streptomycin and 1% non-essential amino acids. HEK-293T cells were cultured in high-glucose-formulated DMEM media with 10% FBS, 1% penicillin/streptomycin and 1% glutamine. Logarithmically growing, mycoplasma-negative cells were utilized for all experiments. Following drug treatments, cells were washed free of the drug(s) prior to the performance of the studies described. MV4-11-MITR and OCI-AML3-MITR cells were generated by culturing MV4-11 or OCI-AML3 cells in their LD_90_ concentration of FHD-286 for 96 hours. Dead cells were removed by Ficoll Hypaque centrifugation. Live cells were washed once with complete media to remove residual Ficoll and cultured in complete media until viability was greater than 90% by trypan blue dye exclusion assessment. This process was repeated for a total of 10 shocks.

### Whole Exome Analysis of MV4-11, MV4-11-MITR, OCI-AML3 and OCI-AML3-MITR cells

Whole exome analysis was performed on MV4-11, MV4-11-MITR, OCI-AML3 and OCI-AML3-MITR cells utilizing Agilent Exome 7 (SureSelect Human All Exon v7). The raw paired-end (PE) reads in FASTQ format were aligned to the human reference genome (hg38) for human DNA-Seq, using BWA alignment software.

### Cell Line Authentication

The cell lines utilized in these studies were authenticated in the Characterized Cell Line Core Facility at M.D. Anderson Cancer Center, Houston TX utilizing STR profiling.

### Primary AML blasts

Patient-derived AML cells samples were obtained with informed consent as part of a clinical protocol approved by the Institutional Review Board of The University of Texas, M.D. Anderson Cancer Center. Normal hematopoietic progenitor cells (HPCs) were obtained from delinked, de-identified cord blood samples. Mononuclear cells were purified by Ficoll Hypaque (Axis Shield, Oslo, Norway) density centrifugation following the manufacturer’s protocol. Mononuclear cells were washed once with sterile 1X PBS and suspended in complete RPMI media containing 20% FBS and counted to determine the number of cells isolated prior to immuno-magnetic selection. CD34+ AML blast progenitor cells were purified by immuno-magnetic beads conjugated with anti-CD34 antibody following the manufacturer’s protocol (StemCell Technologies, Vancouver, British Columbia) prior to utilization in the cell viability assays, RNA expression, and immunoblot analyses.

### Assessment of leukemia cell differentiation

Following treatment with FHD-286, cells were harvested and washed with 1X PBS. Cells were re-suspended in 0.5% BSA/PBS and stained with APC-conjugated anti-CD11b antibody [RRID:AB_398456] or APC-conjugated IgG1 isotype control antibody [RRID:AB_398613] in the dark, at 4°C for 15-20 minutes. Cells were washed with 0.5% BSA/PBS by centrifugation at 125 x g for 5 minutes, and then re-suspended in 0.5% BSA/PBS for analysis by flow cytometry. Cells were assessed in the FL-4 fluorescence channels on a BD Accuri CFLow6 flow cytometer. Differentiation of leukemia cells was also determined by examination of cellular/nuclear morphology. Cells were cytospun onto glass slides at 500 rpm for 5 minutes. The cytospun cells were fixed and stained with a Protocol® HEMA3 stain set (Fisher Scientific, Kalamazoo, MI). Cellular/nuclear morphology was assessed by light microscopy. Two hundred cells were counted in at least 5 different sections of the slide for each condition. The % morphologic differentiation is reported relative to control cells. Each experiment was performed at least twice.

### Assessment of percentage non-viable cells

Following designated treatments (72-96 hours), cultured cell lines or patient-derived (PD) AML blast cells, were washed with 1X PBS, stained with TO-PRO-3 iodide (Cat# T3605, Life Technologies, Carlsbad, CA) and analyzed by flow cytometry on a BD Accuri CFlow-6 flow cytometer (BD Biosciences, San Jose, CA). We used matrix dosing of agents in combinations to allow synergy assessment utilizing the SynergyFinder V2 online web application tool and Delta Synergy scores by ZIP method.

**Methods for Sequencing of primary de novo AML blast cells, ChIP-Seq analysis of epigenetic state in AML cells *in vitro,* Bulk ATAC-Seq analysis, Transcriptome Analysis, Single cell multiomic ATAC and RNA-Seq, SDS-PAGE and immunoblot analyses, Proteomic profiling, Single cell next-generation mass cytometry ‘CyTOF’ analysis of MLL1r and mtNPM1-expressing AML cells and the in vivo mouse models are detailed in the Supplemental Materials and Methods.**

### Data and Software availability

ATAC-Seq, sc-ATAC-Seq, ChIP-Seq, bulk RNA-Seq and sc-RNA-Seq datasets have been deposited in GEO under Accession IDs. The mass spectrometry proteomics data have been deposited to the ProteomeXchange Consortium via the PRIDE partner repository with the dataset identifier “PXDxxxxxx”

## Acknowledgements

The authors would like to thank the Advanced Technology Genomics Core (ATGC), Flow Cytometry and Cellular Imaging (FCCI) Core Facility, which are supported by the MD Anderson Cancer Center Support Grant 5P30 CA016672-40. NextGen sequencing studies performed utilizing the NovaSeq6000 were supported by a grant from the NIH (1S10OD024977-01). Single cell multiomics analyses on patient-derived AML cells were supported by an Epigenetic Services Grant Program Award from Active Motif (W.F.). The BCM Mass Spectrometry Proteomics Core is supported by the Dan L. Duncan Comprehensive Cancer Center Award (P30 CA125123), CPRIT Core Facility Awards (RP170005 and RP210227), Intellectual Developmental Disabilities Research Center Award (P50 HD103555), and NIH High End Instrument Award (S10 OD026804, Orbitrap Exploris 480). K. N. B. was supported by a grant from the N.I.H (R01 CA255721). This research is supported in part by the M.D. Anderson Cancer Center Leukemia SPORE (P50 CA100632).

## Author Contributions

K. N. B. designed the study, analyzed data, and wrote the manuscript. X. R. and X. S. performed bioinformatics analyses. W. F., C. P. M., C. E. B., K. D., J. A. D., H. H., S. L. and J. W. performed research and analyzed the data. A. J. and A. M. performed the mass spectrometry analyses and analyzed the data. J. P., M. C., M. H., B. C., T. M. K., N. D., K. S., K. T., D. H., P. R., R. S. L. B. F. and C. D. D. contributed critical reagents. W. F. also wrote the manuscript.

