## Supplemental Figure Legends for "BRG1/BRM inhibitor targets AML stem cells and exerts superior preclinical efficacy combined with BET or Menin inhibitor"

**Supplemental Figure 1. SMARCA4 is a dependency in AML cells and treatment with FHD-286 induces differentiation of AML cells and decreases cell viability of PD, mtTP53 expressing AML cells while sparing normal CD34+ HPCs. A.** CRISPR-dependency (Chronos) screening scores from DepMap show increased dependency of AML cells on SMARCA4. The lower the score, the greater the dependency of the AML cell line on SMARCA4. **B**. RNAi dependency screen (DEMETER2) from DepMap shows that BRG1 knockdown by shRNA also represents a dependency in AML cells. **C**. Immunoblot analysis of BRG1 (SMARCA4) and BRM (SMARCA2), in sixteen PD, mtNPM1 and MLL1-rearranged AML cells. The expression levels of β-Actin in the cell lysates served as the loading control. **D.** MOLM13, OCI-AML3 and MV4-11 cells were treated with 30 nM of FHD-286 for 7 days. Following this, cells were assessed for the mean fluorescent intensity (MFI) of CD11b by flow cytometry. Columns, mean of three experiments, Bars, S.E.M. **E**. MOLM13, OCI-AML3 and MV4-11 cells were treated with 30 nM of FHD-286 for 7 days. At the end of treatment, cells were cytospun onto glass slides and assessed for morphologic differentiation. Representative light microscopy images of differentiated cells are shown. **F**. Representative light microscopy images of short-term-cultured PD AML cells treated for 7 days with 30 nM of FHD-286. At the end of treatment, cells were cytospun onto glass slides and stained with H & E or FACS sorted for CD11b expression and cultured for an additional 7 days prior to cytospin and H & E staining. **G**. Normal CD34+ HPCs (n=5) were treated with the indicated concentrations of FHD-286 for 72 hours. At the end of treatment, cells were stained with TO-PRO-3 iodide and the % non-viable cells were determined by flow cytometry. **H**. Oncoplot of NGS-determined mutations in patient-derived MLL1 rearranged and mtNPM1-expressing AML cells. **I**. Mutant TP53-expressing AML MOLM13 TP53-R175H and MOLM13 TP53-R248Q cells were treated with the indicated concentrations of FHD-286 for 96 hours. Then, cells were stained with TO-PRO-3 iodide and the % non-viable cells were determined by flow cytometry. Columns, mean of three experiments, Bars, S.E.M. **J-K.** MV4-11, OCI-AML3 and their drug tolerant/resistant counterparts MV4-11-MITR and OCI-AML3-MITR were treated with the indicated concentrations of SNDX-50469 for 96 hours. Following this, cells were stained with TO-PRO-3 iodide and the % non-viable cells were determined by flow cytometry. Columns, mean of three experiments, Bars, S.E.M. **L**. Genomic DNA was isolated from MV4-11-MITR and OCI-AML3-MITR cells and analyzed for the presence of mutations in Menin utilizing custom-designed TaqMan SNP probes specific for each mutation site.

**Supplemental Figure 2. Treatment with FHD-286 depletes H3K27Ac occupancy on chromatin, reduces super enhancer activity and alters core regulatory circuitry in AML cells. A.** MOLM13 cells were treated with 100 nM of FHD-286 for 16 hours. ChIP Seq analysis was conducted with anti-H3K27Ac antibody. Panel shows the genome-wide peak profile and heat map of H3K27Ac occupancy at peak center +/- 5 kb resolution. **B**. Core regulatory circuitry analysis was conducted on H3K27Ac ChIP-Seq data from control and FHD-286-treated MOLM13 cells. The panel shows the CRC score and total number of transcription factors in the CRC of untreated and FHD-286-treated cells.

**Supplemental Figure 3. RNA-Seq analysis of FHD286-treated MOLM13 cells demonstrates significant reduction in the normalized enrichment scores for MYC, mTORC1, E2F, Interferon-gamma, IL6-JAK-STAT3, as well as of inflammatory response and oxidative phosphorylation genes. A.** MOLM13 cells were treated in biologic triplicates with the indicated concentration of FHD-286 for 16 hours. Total RNA was isolated and RNA-Seq analysis was conducted. The heat map indicates the number of mRNAs induced or depleted at a $\geq$1.25 fold-change and $p < 0.05$. **B**. Gene set enrichment analysis of FHD-286-treated MOLM13 cells compared to REACTOME pathways. Normalized enrichment scores are shown. All q-values are less than 0.1. **C.** Log2 fold-change in MYC_TARGETS_V1 and V2 genes in FHD-286-treated compared to control MOLM13 cells. **D-F**. Log2 fold-change in known isoforms of MYC, SPI1 and BCL2 in FHD-286-treated MOLM13 cells. **G-H**. Heat map and log2 fold-change in endogenous retroviral elements in FHD-286-treated compared to control MOLM13 cells.

**Supplemental Figure 4. Treatment with FHD-286 reduces chromatin accessibility and depletes transcription factor binding motif enrichment in the CMP cluster of patient-derived mtNPM1 + FLT3-ITD AML cells. A**. Transcription Factor binding motifs decreased in loci with reduced Tn5-transposase accessible chromatin in CMP cluster cells following treatment with FHD-286. The motif name, canonical binding motif and the p-value are shown. **B**. IGV plot of the SPI1 locus in PD mtNPM1 + FLT3-ITD AML CMP cluster cells with and without FHD-286 treatment. **C**. Patient-derived MLL-AF9 expressing AML cells from a bone marrow aspirate were treated with 100 nM of FHD-286 for 16 hours. Multiomics (combined sc-ATAC-Seq and sc-RNA-Seq) analyses were performed on isolated nuclei. The UMAP plot shows the SingleR-determined composition of the individual cell clusters in the control and FHD-286-treated cells. **D**. Percentage of each cell type in the clusters of cells from control and FHD-286 treated MLL-AF9 expressing AML cells. Arrows indicate clusters with increased or decreased numbers of cells in the FHD-286-treated sample compared to the control sample. **E**. Gene set enrichment analysis of FHD-286-treated cells over control cells. All q-values are less than 0.1. **F**. Volcano plot of sc-RNA-Seq expression changes ($\geq$1.25 fold change and $p < 0.05$) in the CMP cluster following treatment with FHD-286 compared to control MLL-AF9 expressing AML cells.

**Supplemental Figure 5. FHD-286 mediated protein expression alterations include c-Myc, PU.1 and CDK6 in phenotypically defined AML cells with MLLr or mtNPM1. A**. MOLM13 and PD mtNPM1 expressing AML cells were treated with 100 nM of FHD-286 for 48 hours. Total proteome profiling was conducted by mass spectrometry analysis. The tables show the overlapping depleted (75 proteins) and induced (52 proteins) protein expressions with a fold change greater than 1.25 and a p value <0.05. **B**. Log2 fold-change of induced protein expressions in FHD-286-treated MOLM13 and PD mtNPM1-expressing AML cells compared to REACTOME pathways. **C**. Volcano plot (log2 fold-change versus –log10 p-value) of mass spectrometry-determined protein expression changes in FHD-286-treated compared to control MOLM13 cells. **D**. Heat map of protein expression changes in control and FHD-286-treated MOLM13 cells compared to HALLMARK_MYC_TARGETS_V2 and HALLMARK_INFLAMMATORY_RESPONSE pathway datasets. **E**. Volcano plot (log2 fold-change versus –log10 p-value) of mass spectrometry-determined protein expression changes in FHD-286-treated compared to control PD mtNPM1-expressing AML cells. **F**. Enrichment plots of FHD-286-treated PD mtNPM1-expressing AML cells compared to HALLMARK_MYC_TARGETS_V2, HALLMARK_E2F_TARGETS and HALLMARK_G2M_CHECKPOINT pathway datasets. **G.** MOLM13 cells were treated with 100 nM of FHD-286 for the indicated times. Total cell lysates were prepared and immunoblot analyses were conducted. The expression levels of GAPDH served as the loading control.

**Supplemental Figure 6. FHD-286 exhibits in vivo efficacy against AML initiating stem cells and aggressive mtNPM1 + FLT3-ITD + F691L AML PDX models. A**. Patient-derived mtNPM1 + FLT3-ITD Luc/GFP cells (AML#1 in the oncoplot) were ex vivo treated with 10 and 30 nM of FHD-286 for 96 hours. Following this, equal numbers of cells (2e6 cells/mouse) were tail vein infused into pre-irradiated (2.5Gy) NSG mice (n=5 per cohort). Mice were monitored daily for symptoms of acute leukemia. Luciferase signal in the mice was determined by IVIS (Xenogen) imaging at four weeks post-infusion of the AML cells. Panel shows representative bioluminescent images of mice from each ex vivo treatment cohort. **B**. Kaplan-Meier survival curve of NSG mice infused with equal numbers of patient-derived mtNPM1 + FLT3-ITD Luc/GFP cells and treated with vehicle or 1.5 mg/kg of FHD-286 for 5 weeks. Significance between cohorts was determined by a Mantel-Cox log-rank test. P-values less than 0.05 were considered significant. **C**. Kaplan-Meier survival curve of NSG mice infused with equal numbers of patient-derived mtNPM1 + FLT3-ITD + FLT3-F691L Luc/GFP cells (AML#25 in the oncoplot) and treated with vehicle or 1.5 mg/kg of FHD-286 for 5 weeks. Significance between cohorts was determined by a Mantel-Cox log-rank test. P-values less than 0.05 were considered significant. **D-E**. C57/BL6 mice were treated with vehicle or 1.5 mg/kg of FHD-286 for two weeks. Weights of the mice were monitored on a weekly basis. Peripheral blood was collected by retro-orbital bleed and complete blood count (CBC) analyses were performed. Following this, the mice were allowed to recover for two weeks with no treatment and CBCs were repeated. ** =p <0.01 for platelets in FHD-286 treated mice compared to vehicle treated mice.

**Supplemental Figure 7. Co-treatment with FHD-286 and decitabine induces synergistic *in vitro* lethality in cultured cell lines with MLL-r and mtNPM1 while treatment with FHD-286 and OTX015 or Menin inhibitor SNDX-50469 is relatively sparing of normal CD34+ HPCs. A**. MOLM13, MV4-11 and OCI-AML3 cells were treated with FHD-286 (dose range: 10 -50 nM) and decitabine (dose range: 50-250 nM) for 96 hours. The % non-viable cells was determined by TO-PRO-3 iodide staining and flow cytometry. Delta Synergy scores were calculated by the ZIP method utilizing the SynergyFinder web application. The table shows the mean Delta Synergy score for the combinations in the three AML cell lines. **B-C**. Normal CD34+ HPCs (n=3) were treated with the indicated concentrations of FHD-286 and/or OTX015 or SNDX-50469 for 72 hours. The % non-viable cells was determined by TO-PRO-3 iodide staining and flow cytometry. **D**. PD MLL-AF9 + FLT3-TKD Luc/GFP AML cells were infused into pre-irradiated (2.5Gy) NSG mice and monitored for 5 days. Treatment was initiated when leukemia engraftment was documented by bioluminescent imaging. Mice were treated with vehicle, 1.5 mg/kg of FHD-286, 1 mg/kg of decitabine, 30 mg/kg of OTX015, FHD-286 plus decitabine, or FHD-286 plus OTX015 for 3 weeks. Total bioluminescent flux was determined for each cohort of mice. The panel shows representative bioluminescent images for the FHD-286 and/or OTX015 treated cohorts of mice.

**Supplemental Table 1. NGS-determined mutations and their associated variant allele frequency (VAF) in the PDX AML cells utilized.**
