## Supplemental Figures for "BRG1/BRM inhibitor targets AML stem cells and exerts superior preclinical efficacy combined with BET or Menin inhibitor"

Figure S1

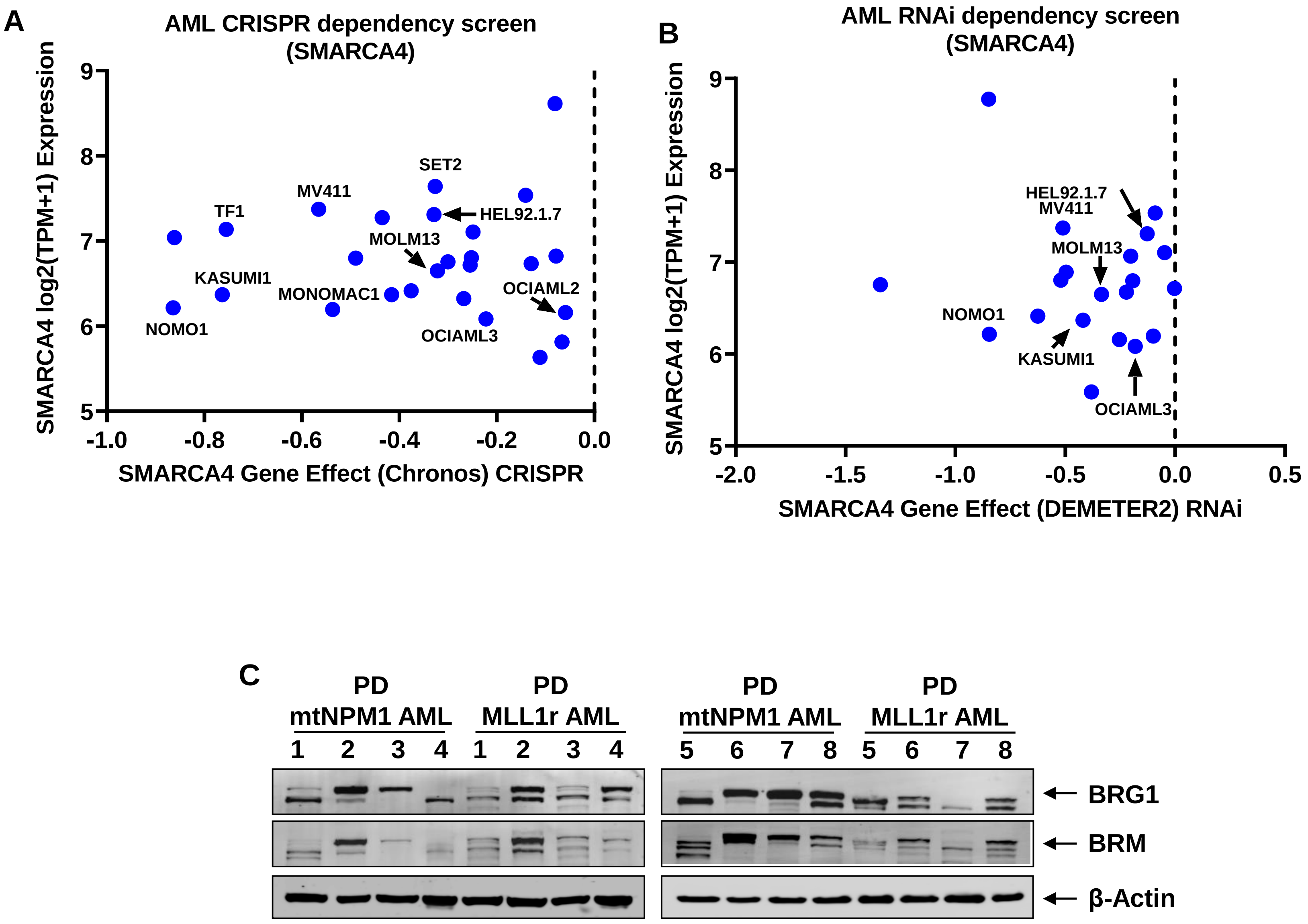
A
AML CRISPR dependency screen (SMARCA4)
SMARCA4 log2(TPM+1) Expression
SMARCA4 Gene Effect (Chronos) CRISPR
SET2
MV411
TF1
HEL92.1.7
MOLM13
KASUMI1
MONOMAC1
OCIAML2
NOMO1
OCIAML3
B
AML RNAi dependency screen (SMARCA4)
SMARCA4 log2(TPM+1) Expression
SMARCA4 Gene Effect (DEMETER2) RNAi
HEL92.1.7
MV411
MOLM13
NOMO1
KASUMI1
OCIAML3
C
PD mtNPM1 AML
PD MLL1r AML
1 2 3 4 1 2 3 4
5 6 7 8 5 6 7 8
BRG1
BRM
β-Actin

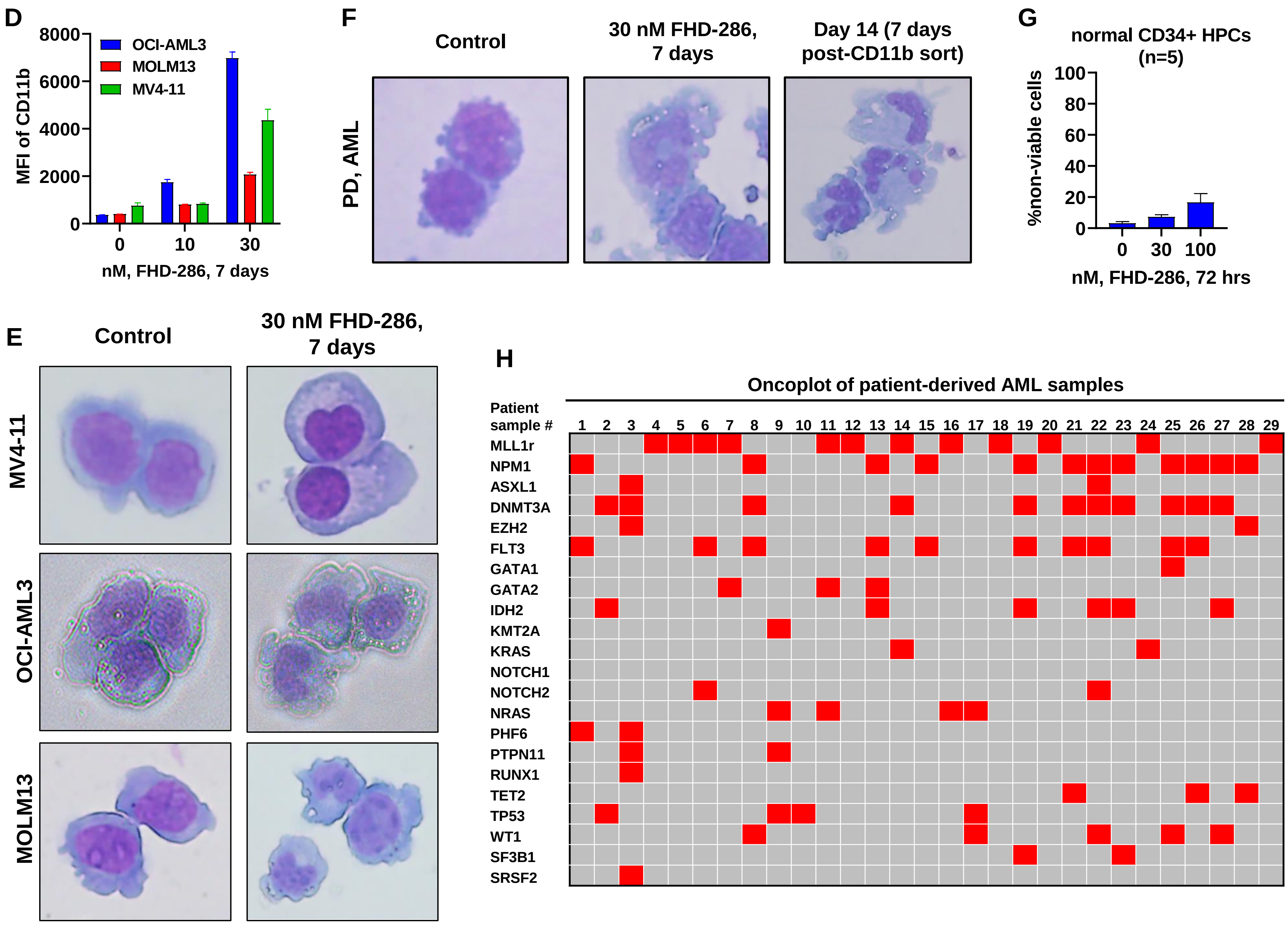
D
MFI of CD11b
8000
6000
4000
2000
0
OCI-AML3
MOLM13
MV4-11
0
10
30
nM, FHD-286, 7 days
F
Control
30 nM FHD-286, 7 days
Day 14 (7 days post-CD11b sort)
PD, AML
G
normal CD34+ HPCs (n=5)
%non-viable cells
100
80
60
40
20
0
0
30
100
nM, FHD-286, 72 hrs
E
Control
30 nM FHD-286, 7 days
MV4-11
OCI-AML3
MOLM13
H
Oncoplot of patient-derived AML samples
Patient sample #
1 2 3 4 5 6 7 8 9 10 11 12 13 14 15 16 17 18 19 20 21 22 23 24 25 26 27 28 29
MLL1r
NPM1
ASXL1
DNMT3A
EZH2
FLT3
GATA1
GATA2
IDH2
KMT2A
KRAS
NOTCH1
NOTCH2
NRAS
PHF6
PTPN11
RUNX1
TET2
TP53
WT1
SF3B1
SRSF2

Figure S1

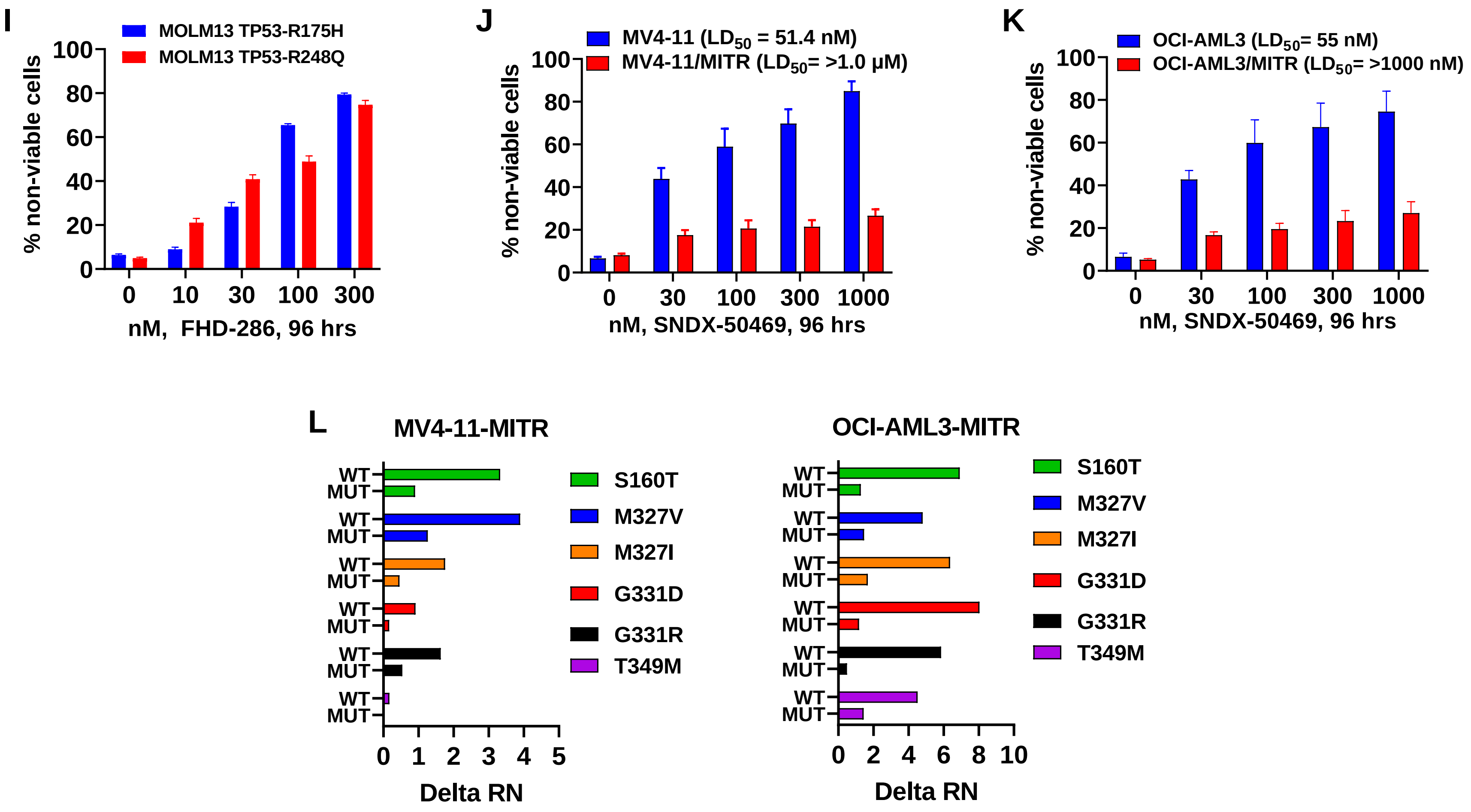
I
MOLM13 TP53-R175H
MOLM13 TP53-R248Q
% non-viable cells
nM, FHD-286, 96 hrs
J
MV4-11 ($LD_{50}$ = 51.4 nM)
MV4-11/MITR ($LD_{50}$= >1.0 μM)
% non-viable cells
nM, SNDX-50469, 96 hrs
K
OCI-AML3 ($LD_{50}$= 55 nM)
OCI-AML3/MITR ($LD_{50}$= >1000 nM)
% non-viable cells
nM, SNDX-50469, 96 hrs
L
MV4-11-MITR
OCI-AML3-MITR
WT
MUT
S160T
M327V
M327I
G331D
G331R
T349M
Delta RN

Figure S1

### Figure S2

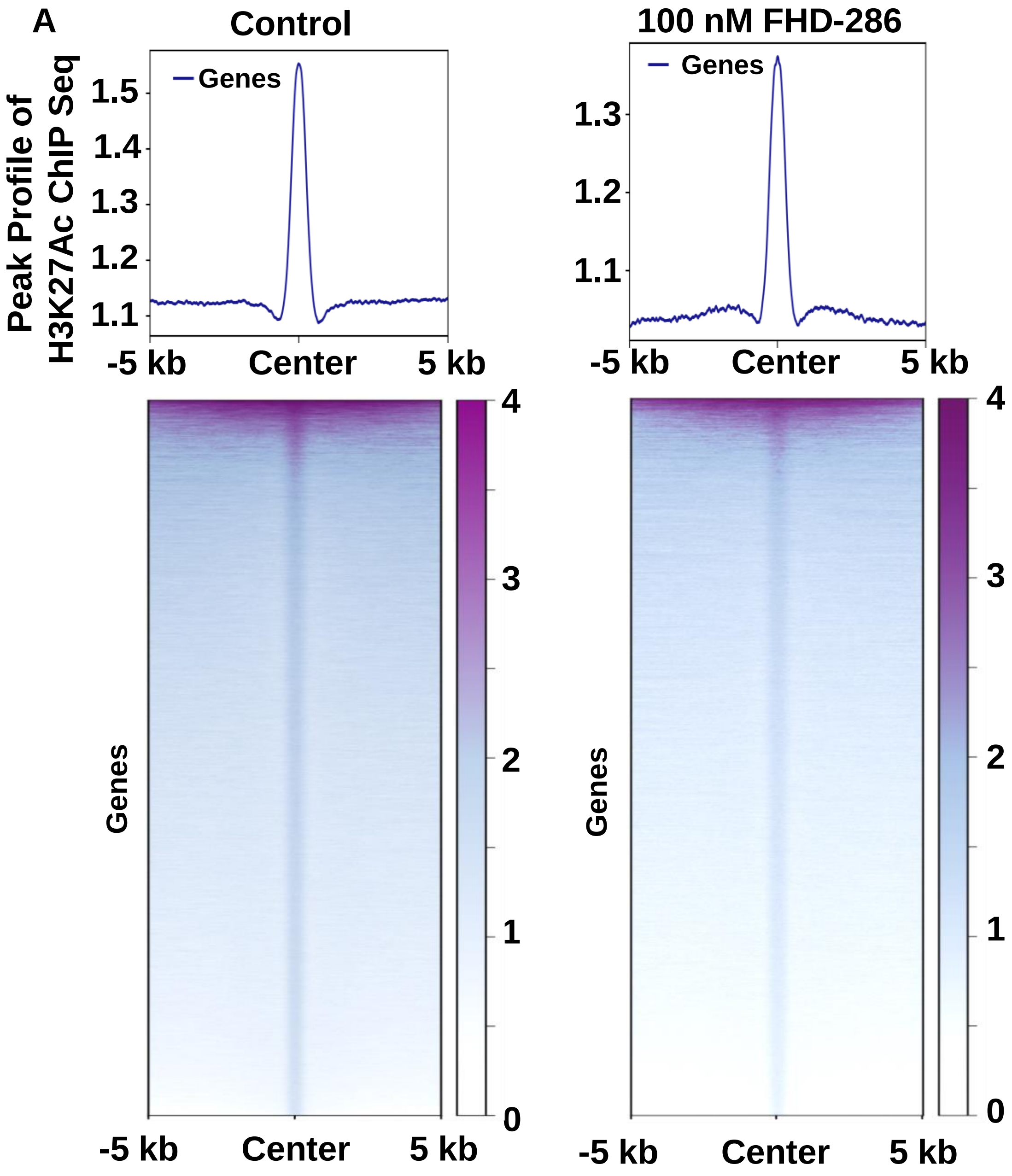
A
Control
100 nM FHD-286
Peak Profile of
H3K27Ac ChIP Seq
Genes
1.5
1.4
1.3
1.2
1.1
-5 kb
Center
5 kb
4
3
2
1
0

B

### Core Transcriptional Regulatory Circuitry (CRC) in MOLM13 cells

#### Control

Scores:5.81 TFs Num:47

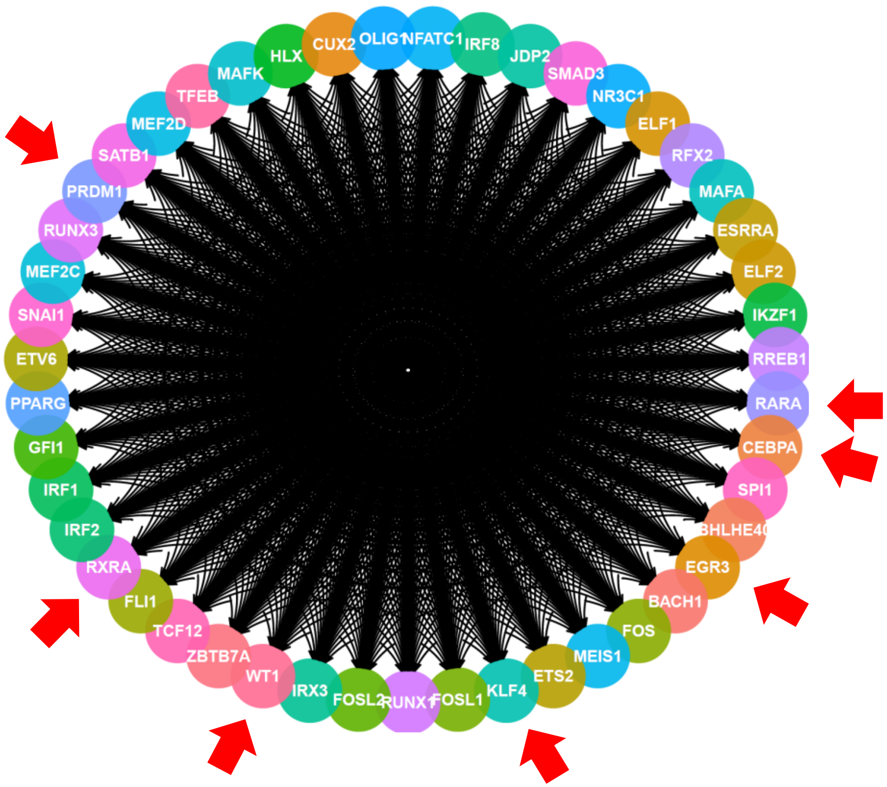


#### 100 nM FHD-286

Scores:4.71 TFs Num:31

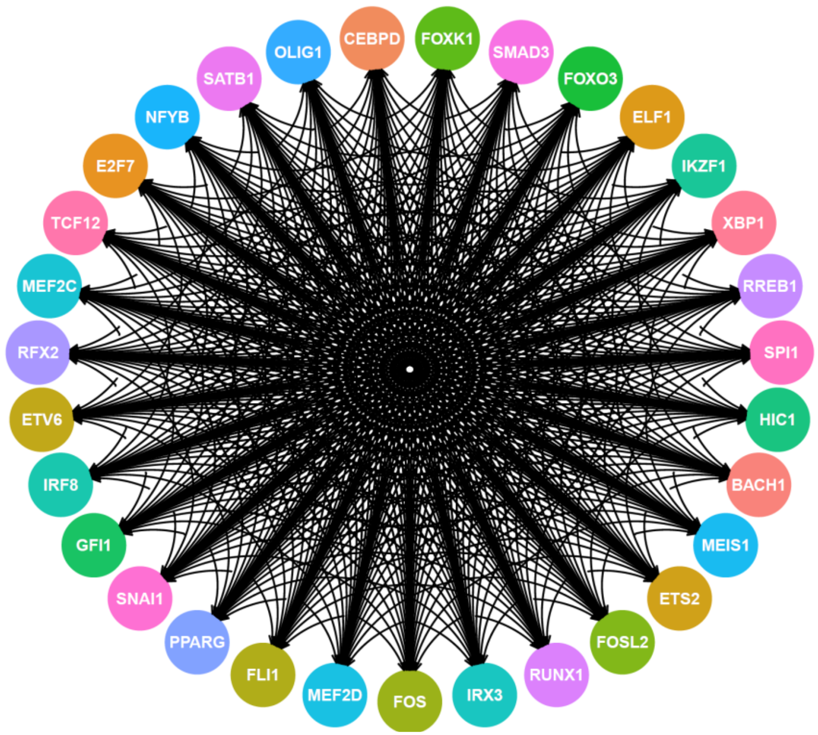

Figure S3

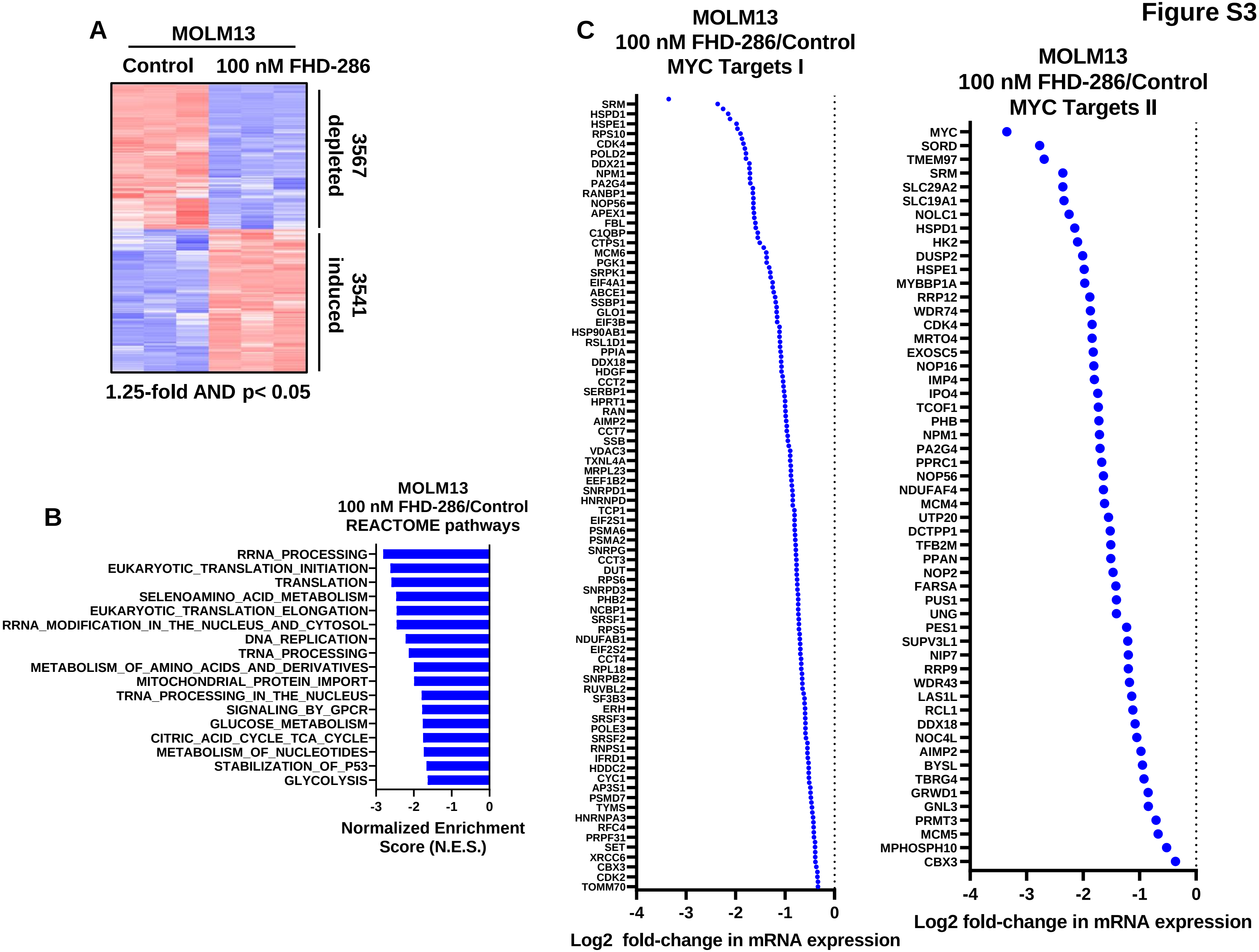

A
MOLM13
Control
100 nM FHD-286
3567 depleted
3541 induced
1.25-fold AND p< 0.05
B
MOLM13
100 nM FHD-286/Control
REACTOME pathways
RRNA_PROCESSING
EUKARYOTIC_TRANSLATION_INITIATION
TRANSLATION
SELENOAMINO_ACID_METABOLISM
EUKARYOTIC_TRANSLATION_ELONGATION
RRNA_MODIFICATION_IN_THE_NUCLEUS_AND_CYTOSOL
DNA_REPLICATION
TRNA_PROCESSING
METABOLISM_OF_AMINO_ACIDS_AND_DERIVATIVES
MITOCHONDRIAL_PROTEIN_IMPORT
TRNA_PROCESSING_IN_THE_NUCLEUS
SIGNALING_BY_GPCR
GLUCOSE_METABOLISM
CITRIC_ACID_CYCLE_TCA_CYCLE
METABOLISM_OF_NUCLEOTIDES
STABILIZATION_OF_P53
GLYCOLYSIS
-3 -2 -1 0
Normalized Enrichment Score (N.E.S.)
C
MOLM13
100 nM FHD-286/Control
MYC Targets I
SRM
HSPD1
HSPE1
RPS10
CDK4
POLD2
DDX21
NPM1
PA2G4
RANBP1
NOP56
APEX1
FBL
C1QBP
CTPS1
MCM6
PGK1
SRPK1
EIF4A1
ABCE1
SSBP1
GLO1
EIF3B
HSP90AB1
RSL1D1
PPIA
DDX18
HDGF
CCT2
SERBP1
HPRT1
RAN
AIMP2
CCT7
SSB
VDAC3
TXNL4A
MRPL23
EEF1B2
SNRPD1
HNRNPD
TCP1
EIF2S1
PSMA6
PSMA2
SNRPG
CCT3
DUT
RPS6
SNRPD3
PHB2
NCBP1
SRSF1
RPS5
NDUFAB1
EIF2S2
CCT4
RPL18
SNRPB2
RUVBL2
SF3B3
ERH
SRSF3
POLE3
SRSF2
RNPS1
IFRD1
HDDC2
CYC1
AP3S1
PSMD7
TYMS
HNRNPA3
RFC4
PRPF31
SET
XRCC6
CBX3
CDK2
TOMM70
-4 -3 -2 -1 0
Log2 fold-change in mRNA expression
MOLM13
100 nM FHD-286/Control
MYC Targets II
MYC
SORD
TMEM97
SRM
SLC29A2
SLC19A1
NOLC1
HSPD1
HK2
DUSP2
HSPE1
MYBBP1A
RRP12
WDR74
CDK4
MRTO4
EXOSC5
NOP16
IMP4
IPO4
TCOF1
PHB
NPM1
PA2G4
PPRC1
NOP56
NDUFAF4
MCM4
UTP20
DCTPP1
TFB2M
PPAN
NOP2
FARSA
PUS1
UNG
PES1
SUPV3L1
NIP7
RRP9
WDR43
LAS1L
RCL1
DDX18
NOC4L
AIMP2
BYSL
TBRG4
GRWD1
GNL3
PRMT3
MCM5
MPHOSPH10
CBX3
-4 -3 -2 -1 0
Log2 fold-change in mRNA expression

Figure S3

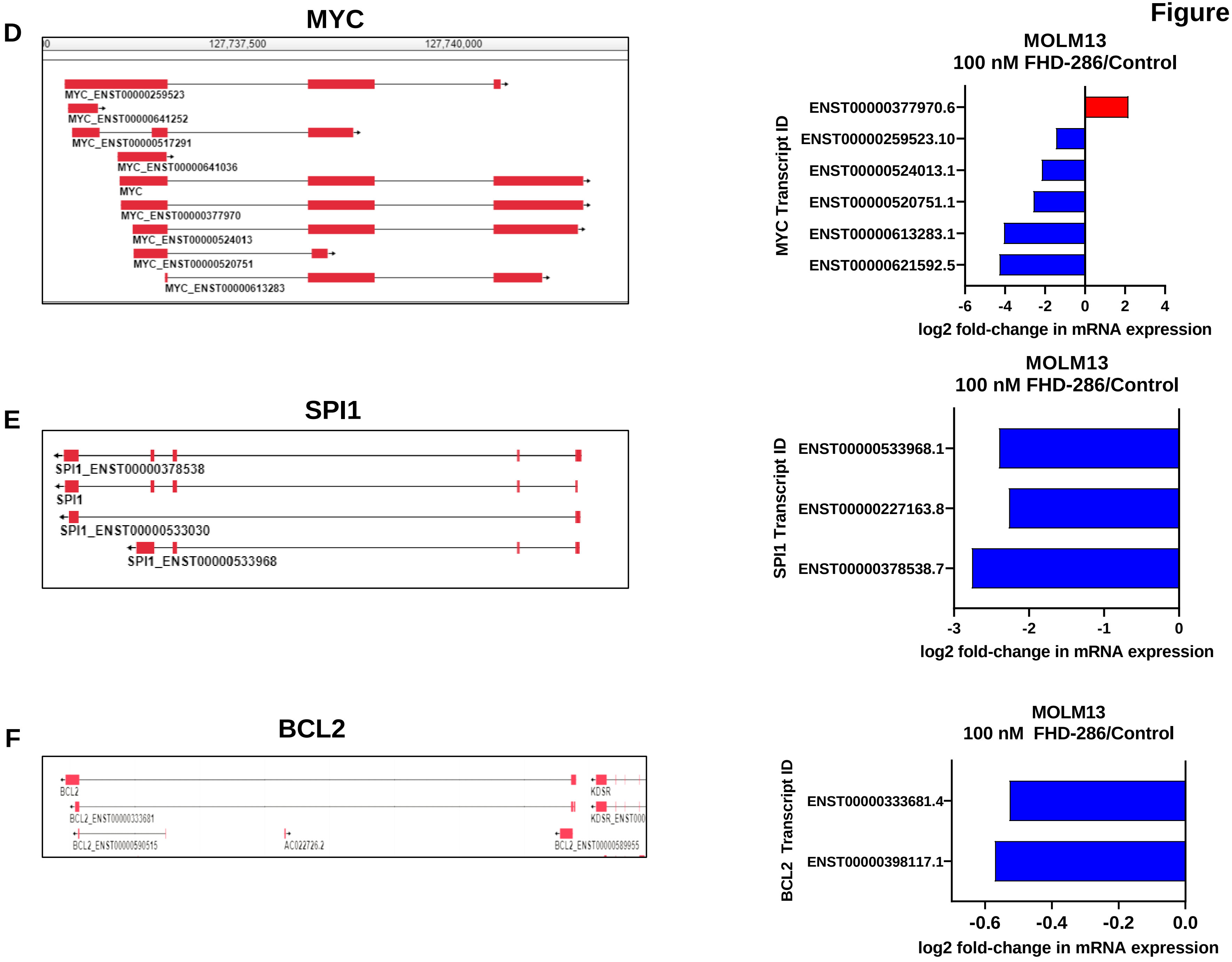
D
MYC
127,737,500
127,740,000
MYC_ENST00000259523
MYC_ENST00000641252
MYC_ENST00000517291
MYC_ENST00000641036
MYC
MYC_ENST00000377970
MYC_ENST00000524013
MYC_ENST00000520751
MYC_ENST00000613283
MOLM13
100 nM FHD-286/Control
MYC Transcript ID
ENST00000377970.6
ENST00000259523.10
ENST00000524013.1
ENST00000520751.1
ENST00000613283.1
ENST00000621592.5
-6
-4
-2
0
2
4
log2 fold-change in mRNA expression
E
SPI1
SPI1_ENST00000378538
SPI1
SPI1_ENST00000533030
SPI1_ENST00000533968
MOLM13
100 nM FHD-286/Control
SPI1 Transcript ID
ENST00000533968.1
ENST00000227163.8
ENST00000378538.7
-3
-2
-1
0
log2 fold-change in mRNA expression
F
BCL2
BCL2
BCL2_ENST00000333681
BCL2_ENST00000590515
AC022726.2
BCL2_ENST00000589955
KDSR
MOLM13
100 nM FHD-286/Control
BCL2 Transcript ID
ENST00000333681.4
ENST00000398117.1
-0.6
-0.4
-0.2
0.0
log2 fold-change in mRNA expression

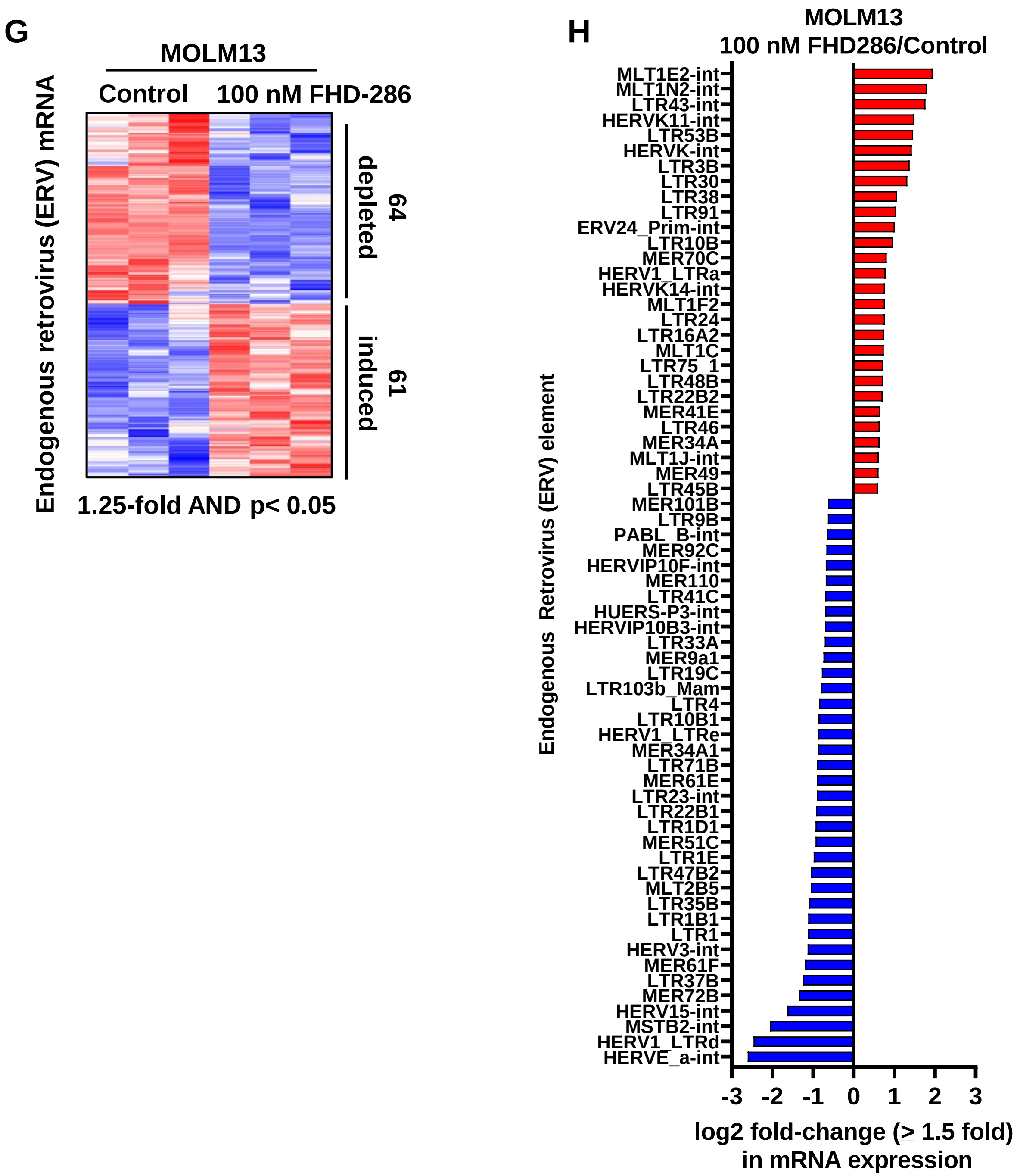
G
MOLM13
Control
100 nM FHD-286
Endogenous retrovirus (ERV) mRNA
64 depleted
61 induced
1.25-fold AND p< 0.05
H
MOLM13
100 nM FHD286/Control
Endogenous Retrovirus (ERV) element
MLT1E2-int
MLT1N2-int
LTR43-int
HERVK11-int
LTR53B
HERVK-int
LTR3B
LTR30
LTR38
LTR91
ERV24_Prim-int
LTR10B
MER70C
HERV1_LTRa
HERVK14-int
MLT1F2
LTR24
LTR16A2
MLT1C
LTR75_1
LTR48B
LTR22B2
MER41E
LTR46
MER34A
MLT1J-int
MER49
LTR45B
MER101B
LTR9B
PABL_B-int
MER92C
HERVIP10F-int
MER110
LTR41C
HUERS-P3-int
HERVIP10B3-int
LTR33A
MER9a1
LTR19C
LTR103b_Mam
LTR4
LTR10B1
HERV1_LTRe
MER34A1
LTR71B
MER61E
LTR23-int
LTR22B1
LTR1D1
MER51C
LTR1E
LTR47B2
MLT2B5
LTR35B
LTR1B1
LTR1
HERV3-int
MER61F
LTR37B
MER72B
HERV15-int
MSTB2-int
HERV1_LTRd
HERVE_a-int
-3 -2 -1 0 1 2 3
log2 fold-change (≥ 1.5 fold) in mRNA expression

**A**

**Decreased TF binding motifs in lost sc-ATAC peaks**

| Motif | Motif Sequence | P-value |
|---|---|---|
| FLI1 | 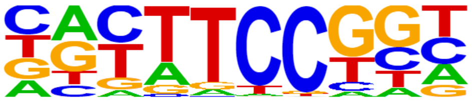 | 1e-39 |
| ETS1 | 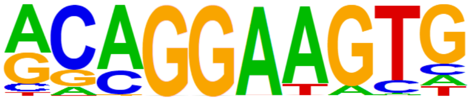 | 1e-38 |
| ERG | 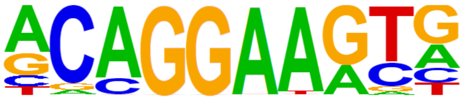 | 1e-35 |
| PU.1 | 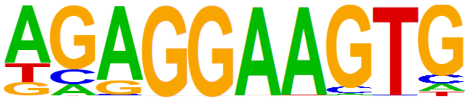 | 1e-33 |
| SPIB | 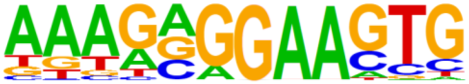 | 1e-29 |
| RUNX1 | 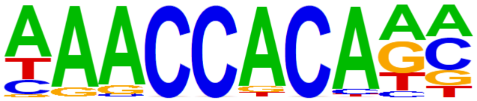 | 1e-16 |

**B**

**sc-ATAC Seq**

**PD mtNPM1 + FLT3-ITD AML**

**Control**

**FHD-286, 16 hrs**

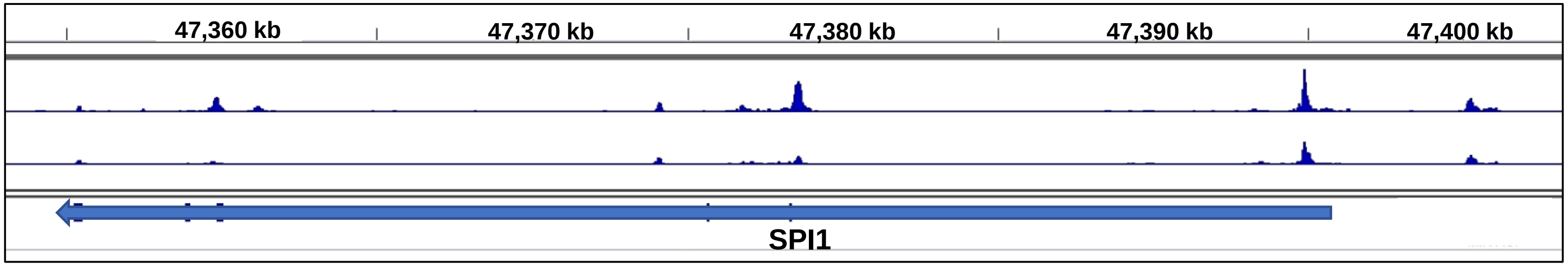

### Figure S4

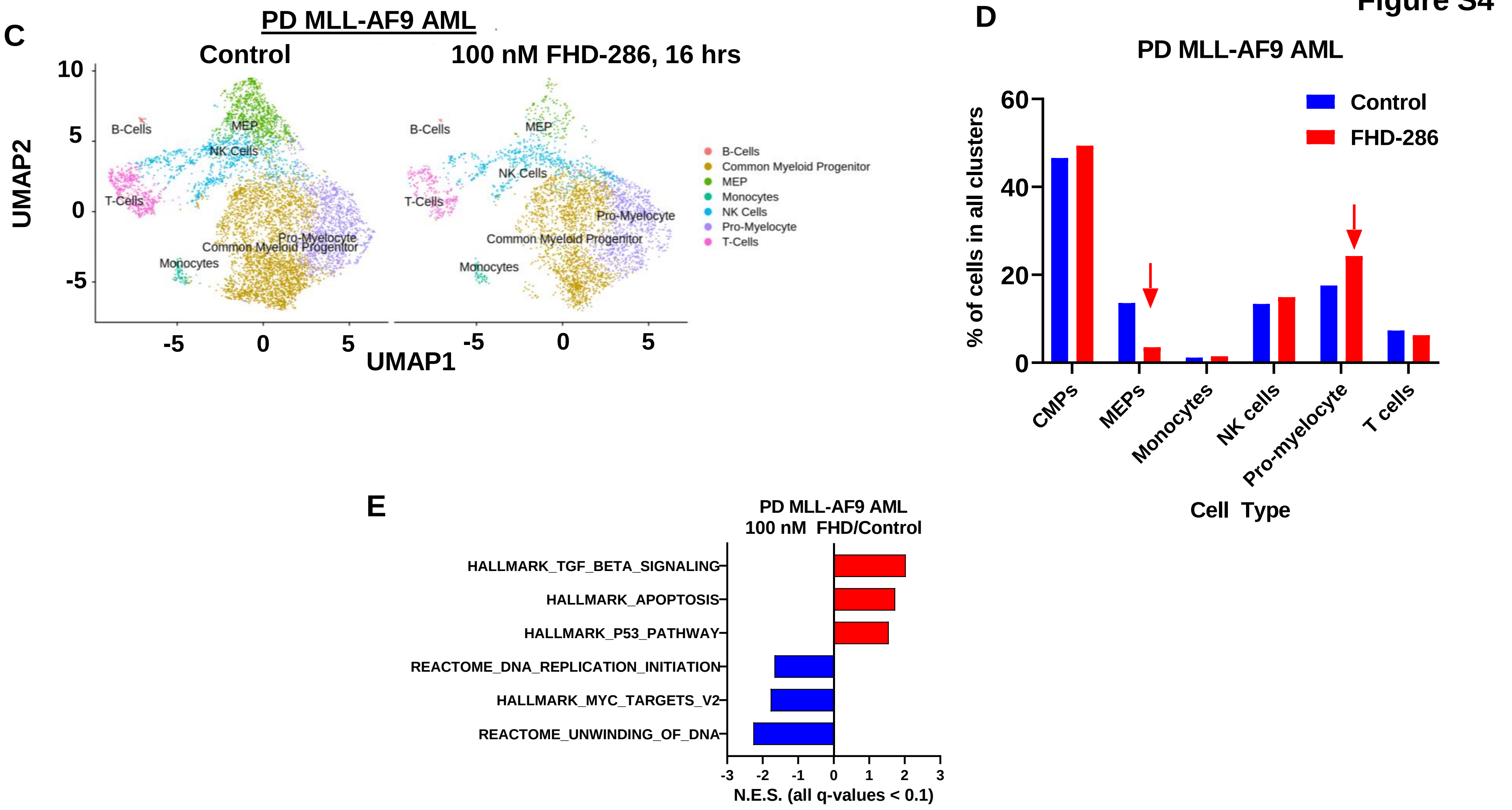
C
PD MLL-AF9 AML
Control
100 nM FHD-286, 16 hrs
UMAP2
UMAP1
10
5
0
-5
-5
0
5
-5
0
5
B-Cells
MEP
NK Cells
T-Cells
Pro-Myelocyte
Common Myeloid Progenitor
Monocytes
B-Cells
Common Myeloid Progenitor
MEP
Monocytes
NK Cells
Pro-Myelocyte
T-Cells
D
PD MLL-AF9 AML
Control
FHD-286
% of cells in all clusters
60
40
20
0
CMPs
MEPs
Monocytes
NK cells
Pro-myelocyte
T cells
Cell Type
E
PD MLL-AF9 AML
100 nM FHD/Control
HALLMARK_TGF_BETA_SIGNALING
HALLMARK_APOPTOSIS
HALLMARK_P53_PATHWAY
REACTOME_DNA_REPLICATION_INITIATION
HALLMARK_MYC_TARGETS_V2
REACTOME_UNWINDING_OF_DNA
-3
-2
-1
0
1
2
3
N.E.S. (all q-values < 0.1)

A

Figure S5

| | | Log2 fold change | |
|---|---|---|---|
| | Protein Name | MOLM13 | PD mtNPM1 AML |
| 1 | COL4A4 | -0.38732 | -2.31531 |
| 2 | FAN1 | -0.67077 | -1.36481 |
| 3 | SEMA4A | -2.30951 | -1.16948 |
| 4 | SPI1 | -0.40088 | -1.0626 |
| 5 | TYMS | -0.53827 | -1.05476 |
| 6 | GPT2 | -0.93609 | -1.0535 |
| 7 | ELANE | -0.45468 | -1.04712 |
| 8 | CST7 | -0.46827 | -1.04643 |
| 9 | CD86 | -0.57047 | -1.03285 |
| 10 | MYC | -0.32393 | -1.01646 |
| 11 | CD180 | -0.96362 | -0.98663 |
| 12 | HLA-DRB5 | -0.43209 | -0.98375 |
| 13 | PRTN3 | -0.77263 | -0.96081 |
| 14 | CDK18 | -0.82492 | -0.93689 |
| 15 | HLA-DRB1 | -0.43209 | -0.93004 |
| 16 | SLC38A5 | -0.86569 | -0.91886 |
| 17 | MATK | -1.01479 | -0.88273 |
| 18 | UNG | -0.53287 | -0.8417 |
| 19 | CD74 | -0.85628 | -0.82798 |
| 20 | SLC43A1 | -1.21766 | -0.81787 |
| 21 | RAD51B | -0.48418 | -0.77519 |
| 22 | CSF2RA | -0.60152 | -0.76262 |
| 23 | EVL | -0.85469 | -0.74353 |
| 24 | VAMP8 | -0.86572 | -0.73438 |
| 25 | TNFRSF10A | -0.41794 | -0.72589 |
| 26 | SLC1A4 | -0.71987 | -0.70746 |
| 27 | HELLS | -0.35324 | -0.70637 |
| 28 | STON1 | -0.62489 | -0.70587 |
| 29 | LGALS9C | -1.31761 | -0.6912 |
| 30 | LGALS9B | -1.31761 | -0.6912 |
| 31 | CCNB1 | -0.45976 | -0.6807 |
| 32 | PRAM1 | -1.20201 | -0.65091 |
| 33 | IL16 | -0.57239 | -0.64596 |
| 34 | TMEM154 | -1.05999 | -0.64403 |
| 35 | LZTS1 | -1.23242 | -0.6382 |
| 36 | ENPP2 | -1.11056 | -0.62413 |
| 37 | S100A4 | -0.46945 | -0.62281 |
| 38 | KIF2C | -0.51748 | -0.62165 |

| | | Log2 fold change | |
|---|---|---|---|
| | Protein Name | MOLM13 | PD mtNPM1 AML |
| 39 | HES1 | -0.93425 | -0.61299 |
| 40 | PRKAR1B | -0.39905 | -0.61061 |
| 41 | SLC38A1 | -0.94892 | -0.60924 |
| 42 | LGALS9 | -0.93792 | -0.60758 |
| 43 | TOP2A | -0.55039 | -0.58554 |
| 44 | APOBR | -0.39854 | -0.57003 |
| 45 | CHEK1 | -0.42911 | -0.53708 |
| 46 | KPNA2 | -0.56797 | -0.52211 |
| 47 | TLR2 | -0.48622 | -0.51065 |
| 48 | MSRA | -0.45713 | -0.50796 |
| 49 | CD44 | -0.52506 | -0.50638 |
| 50 | RRM1 | -0.67575 | -0.5044 |
| 51 | SKAP2 | -0.84166 | -0.4875 |
| 52 | BUB1B | -0.43781 | -0.46589 |
| 53 | CDK4 | -0.43835 | -0.44846 |
| 54 | TFRC | -0.40166 | -0.44837 |
| 55 | FDFT1 | -0.71092 | -0.44708 |
| 56 | RBL1 | -0.38252 | -0.42807 |
| 57 | ITGAL | -0.49285 | -0.42316 |
| 58 | PRIM1 | -0.40613 | -0.40871 |
| 59 | DDX11 | -0.60614 | -0.39564 |
| 60 | ADAM15 | -0.45348 | -0.39065 |
| 61 | SLC29A1 | -0.66308 | -0.38884 |
| 62 | RPS6KA4 | -0.42622 | -0.36851 |
| 63 | CEP55 | -0.37819 | -0.36215 |
| 64 | FYB | -0.48469 | -0.36023 |
| 65 | GRK6 | -0.43822 | -0.35851 |
| 66 | IFRD2 | -0.4405 | -0.3503 |
| 67 | UAP1 | -0.48545 | -0.34636 |
| 68 | SLC19A1 | -0.67034 | -0.34407 |
| 69 | SLC5A6 | -0.47693 | -0.34005 |
| 70 | CTSS | -0.40553 | -0.33548 |
| 71 | MAD2L1 | -0.41576 | -0.32969 |
| 72 | SNAPC4 | -0.43373 | -0.32683 |
| 73 | WIPF1 | -0.46241 | -0.3266 |
| 74 | ICAM2 | -0.60817 | -0.32329 |
| 75 | ITGAE | -0.71223 | -0.32276 |

| | | Log2 fold change | |
|---|---|---|---|
| | Protein Name | MOLM13 | PD mtNPM1 AML |
| 1 | CTTN | 0.855725 | 1.804876 |
| 2 | CA2 | 1.447272 | 1.645852 |
| 3 | MCEMP1 | 2.27595 | 1.395567 |
| 4 | DNAJB2 | 0.420197 | 1.165636 |
| 5 | CDKN1B | 0.955725 | 1.092539 |
| 6 | TDG | 0.355585 | 1.039496 |
| 7 | CLIC3 | 0.539918 | 1.027012 |
| 8 | GNA12 | 0.527192 | 0.945948 |
| 9 | FTH1 | 1.40179 | 0.923542 |
| 10 | IL3RA | 0.801557 | 0.919422 |
| 11 | HMOX1 | 1.828741 | 0.887493 |
| 12 | PSAP | 1.021103 | 0.839591 |
| 13 | DNASE2 | 1.968063 | 0.753455 |
| 14 | HP | 1.270513 | 0.689807 |
| 15 | PTMS | 0.868734 | 0.628843 |
| 16 | PLD3 | 0.667859 | 0.592549 |
| 17 | CLN5 | 0.540784 | 0.582085 |
| 18 | CYLD | 0.593714 | 0.561243 |
| 19 | DEGS1 | 0.387765 | 0.54376 |
| 20 | TFE3 | 0.798033 | 0.501542 |
| 21 | TPM4 | 1.045427 | 0.487587 |
| 22 | SIGLEC6 | 0.861619 | 0.48502 |
| 23 | LRPAP1 | 0.614321 | 0.470576 |
| 24 | NEU1 | 1.027817 | 0.453264 |
| 25 | GALNS | 0.389259 | 0.443153 |
| 26 | ATP6V0A1 | 0.482594 | 0.440274 |

| | | Log2 fold change | |
|---|---|---|---|
| | Protein Name | MOLM13 | PD mtNPM1 AML |
| 27 | CEBPD | 0.435327 | 0.434336 |
| 28 | ARL3 | 0.492295 | 0.432496 |
| 29 | GLG1 | 0.42938 | 0.426285 |
| 30 | CRAT | 0.35467 | 0.422098 |
| 31 | EVI2B | 0.674025 | 0.415998 |
| 32 | VIM | 0.821746 | 0.407409 |
| 33 | GM2A | 0.334251 | 0.405976 |
| 34 | ADGRE5 | 0.490278 | 0.39199 |
| 35 | POR | 0.399704 | 0.391733 |
| 36 | IRS2 | 0.554529 | 0.390232 |
| 37 | ARSB | 0.352165 | 0.386992 |
| 38 | DIAPH1 | 0.398442 | 0.384483 |
| 39 | TMSB4X | 0.642439 | 0.380331 |
| 40 | NPC1 | 0.503186 | 0.361109 |
| 41 | P2RX4 | 0.683568 | 0.358719 |
| 42 | NFKB2 | 0.600754 | 0.349233 |
| 43 | PRKCA | 0.432784 | 0.349223 |
| 44 | GBA | 0.467806 | 0.343543 |
| 45 | CTSB | 1.060807 | 0.342285 |
| 46 | G6PD | 0.611523 | 0.340838 |
| 47 | VTN | 0.589568 | 0.339238 |
| 48 | GAA | 0.514877 | 0.337367 |
| 49 | RAP1B | 0.365911 | 0.337258 |
| 50 | GAB2 | 0.598275 | 0.335548 |
| 51 | FXR2 | 0.362167 | 0.326908 |
| 52 | FPGT | 0.456827 | 0.322182 |

**Figure S5**

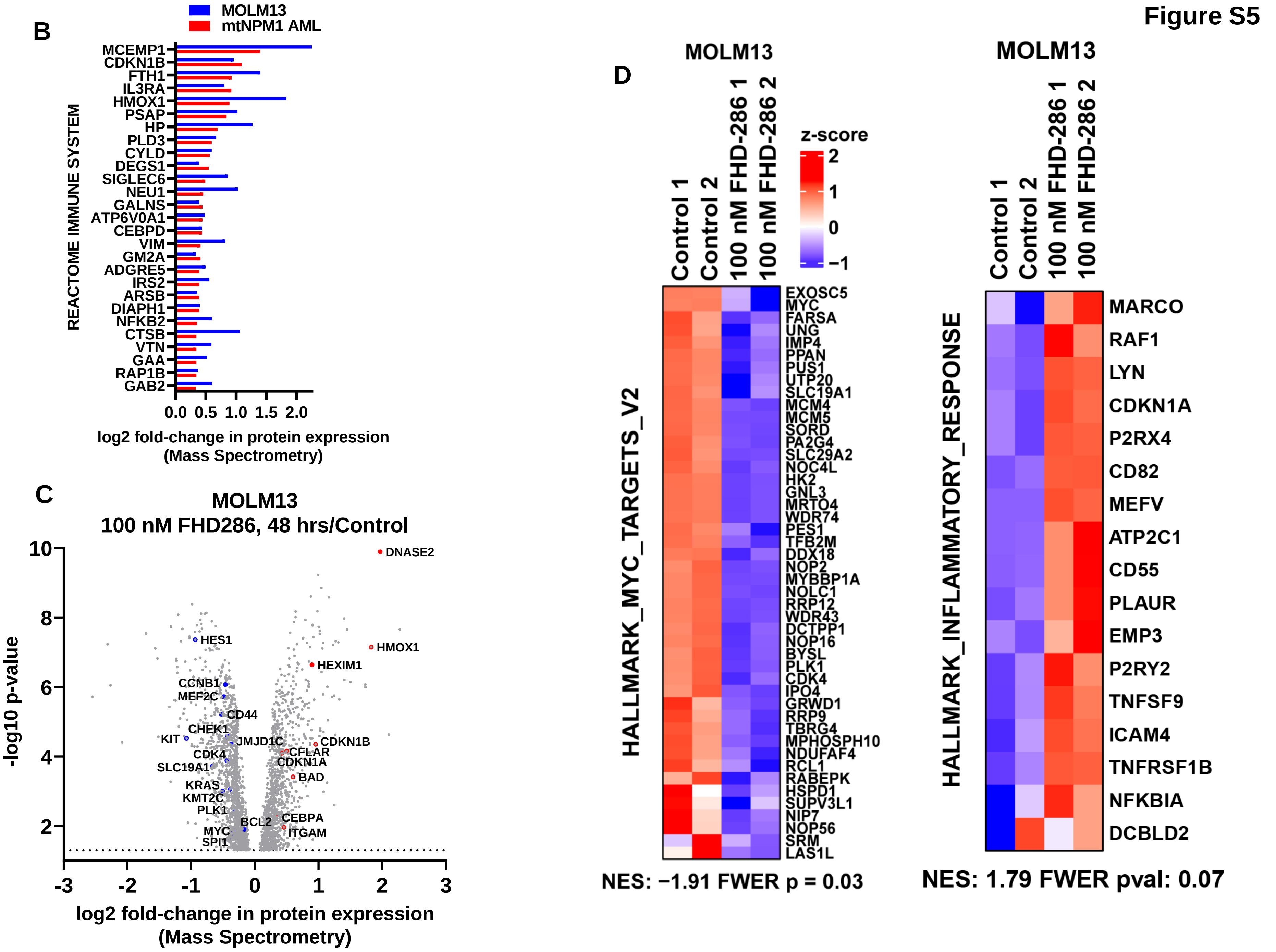
B
MOLM13
mtNPM1 AML
REACTOME IMMUNE SYSTEM
MCEMP1
CDKN1B
FTH1
IL3RA
HMOX1
PSAP
HP
PLD3
CYLD
DEGS1
SIGLEC6
NEU1
GALNS
ATP6V0A1
CEBPD
VIM
GM2A
ADGRE5
IRS2
ARSB
DIAPH1
NFKB2
CTSB
VTN
GAA
RAP1B
GAB2
0.0 0.5 1.0 1.5 2.0
log2 fold-change in protein expression (Mass Spectrometry)
C
MOLM13
100 nM FHD286, 48 hrs/Control
-log10 p-value
DNASE2
HMOX1
HEXIM1
HES1
CCNB1
MEF2C
CD44
CHEK1
KIT
JMJD1C
CDKN1B
CFLAR
CDKN1A
CDK4
SLC19A1
BAD
KRAS
KMT2C
PLK1
BCL2
CEBPA
MYC
ITGAM
SPI1
log2 fold-change in protein expression (Mass Spectrometry)
D
MOLM13
Control 1
Control 2
100 nM FHD-286 1
100 nM FHD-286 2
z-score
HALLMARK_MYC_TARGETS_V2
EXOSC5
MYC
FARSA
UNG
IMP4
PPAN
PUS1
UTP20
SLC19A1
MCM4
MCM5
SORD
PA2G4
SLC29A2
NOC4L
HK2
GNL3
MRTO4
WDR74
PES1
TFB2M
DDX18
NOP2
MYBBP1A
NOLC1
RRP12
WDR43
DCTPP1
NOP16
BYSL
PLK1
CDK4
IPO4
GRWD1
RRP9
TBRG4
MPHOSPH10
NDUFAF4
RCL1
RABEPK
HSPD1
SUPV3L1
NIP7
NOP56
SRM
LAS1L
NES: −1.91 FWER p = 0.03
MOLM13
HALLMARK_INFLAMMATORY_RESPONSE
MARCO
RAF1
LYN
CDKN1A
P2RX4
CD82
MEFV
ATP2C1
CD55
PLAUR
EMP3
P2RY2
TNFSF9
ICAM4
TNFRSF1B
NFKBIA
DCBLD2
NES: 1.79 FWER pval: 0.07

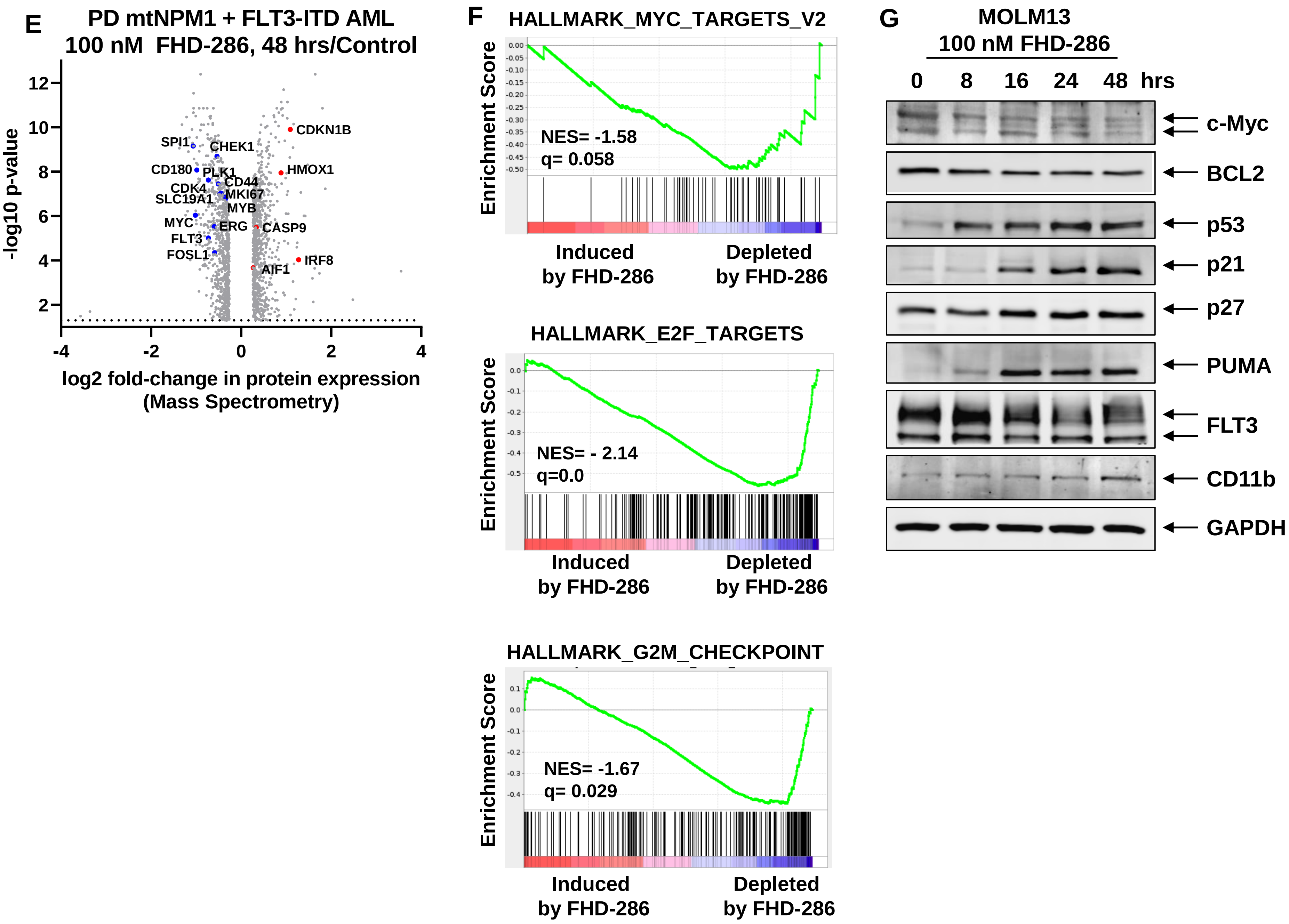

E
PD mtNPM1 + FLT3-ITD AML
100 nM FHD-286, 48 hrs/Control
-log10 p-value
12
10
8
6
4
2
-4
-2
0
2
4
log2 fold-change in protein expression
(Mass Spectrometry)
SPI1
CHEK1
CD180
PLK1
CD44
CDK4
MKI67
SLC19A1
MYB
MYC
ERG
FLT3
FOSL1
CDKN1B
HMOX1
CASP9
IRF8
AIF1
F
HALLMARK_MYC_TARGETS_V2
Enrichment Score
NES= -1.58
q= 0.058
Induced
by FHD-286
Depleted
by FHD-286
HALLMARK_E2F_TARGETS
Enrichment Score
NES= - 2.14
q=0.0
Induced
by FHD-286
Depleted
by FHD-286
HALLMARK_G2M_CHECKPOINT
Enrichment Score
NES= -1.67
q= 0.029
Induced
by FHD-286
Depleted
by FHD-286
G
MOLM13
100 nM FHD-286
0
8
16
24
48
hrs
c-Myc
BCL2
p53
p21
p27
PUMA
FLT3
CD11b
GAPDH


Figure S5

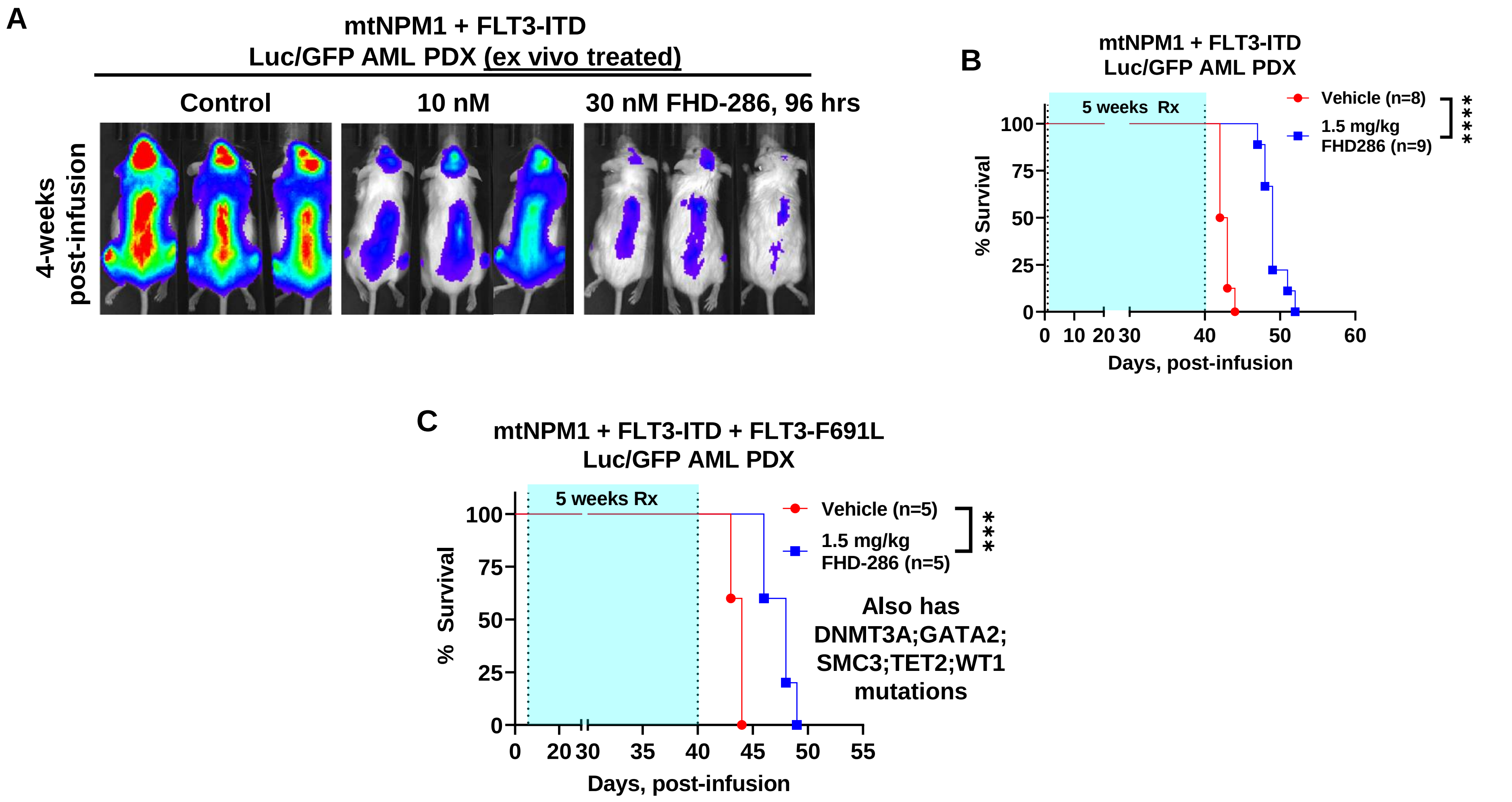
A
mtNPM1 + FLT3-ITD
Luc/GFP AML PDX (ex vivo treated)
Control
10 nM
30 nM FHD-286, 96 hrs
4-weeks
post-infusion
B
mtNPM1 + FLT3-ITD
Luc/GFP AML PDX
5 weeks Rx
Vehicle (n=8)
1.5 mg/kg
FHD286 (n=9)
****
% Survival
100
75
50
25
0
0
10
20
30
40
50
60
Days, post-infusion
C
mtNPM1 + FLT3-ITD + FLT3-F691L
Luc/GFP AML PDX
5 weeks Rx
Vehicle (n=5)
1.5 mg/kg
FHD-286 (n=5)
***
Also has
DNMT3A;GATA2;
SMC3;TET2;WT1
mutations
% Survival
100
75
50
25
0
0
20
30
35
40
45
50
55
Days, post-infusion

Figure S6

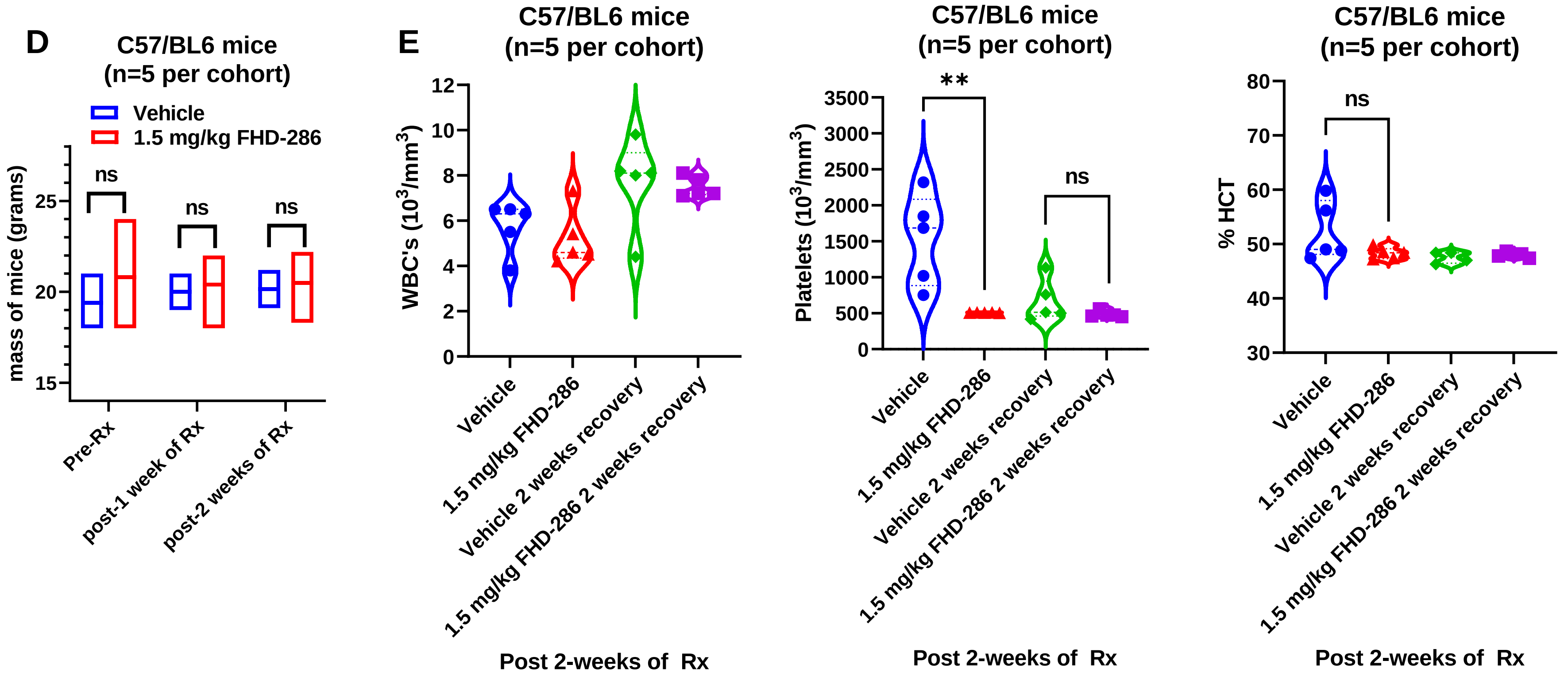
D
C57/BL6 mice
(n=5 per cohort)
Vehicle
1.5 mg/kg FHD-286
mass of mice (grams)
ns
ns
ns
Pre-Rx
post-1 week of Rx
post-2 weeks of Rx
E
C57/BL6 mice
(n=5 per cohort)
WBC's ($10^3$/mm$^3$)
Vehicle
1.5 mg/kg FHD-286
Vehicle 2 weeks recovery
1.5 mg/kg FHD-286 2 weeks recovery
Post 2-weeks of Rx
C57/BL6 mice
(n=5 per cohort)
Platelets ($10^3$/mm$^3$)
**
ns
Post 2-weeks of Rx
C57/BL6 mice
(n=5 per cohort)
% HCT
ns
Post 2-weeks of Rx

Figure S6

Figure S7

A

| | Mean Delta Synergy Score |
|---|---|
| | FHD-286 + decitabine |
| MOLM13 | 42.73 |
| MV4-11 | 34.28 |
| OCI-AML3 | 11.99 |

B

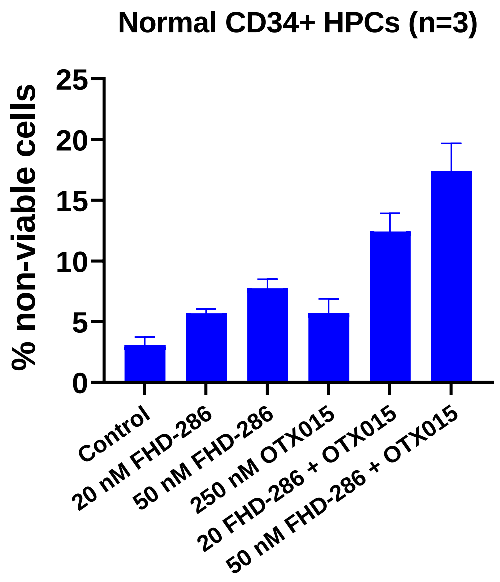


C

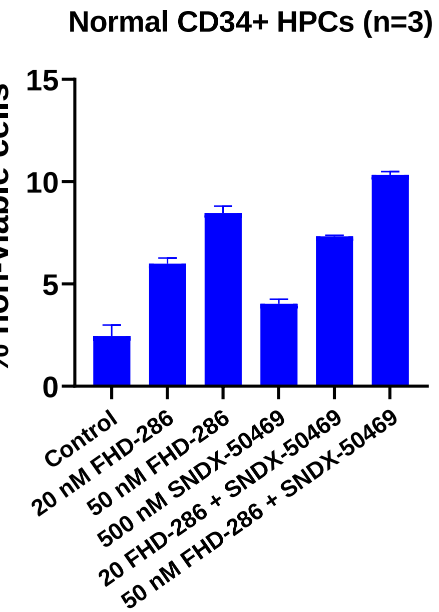


D

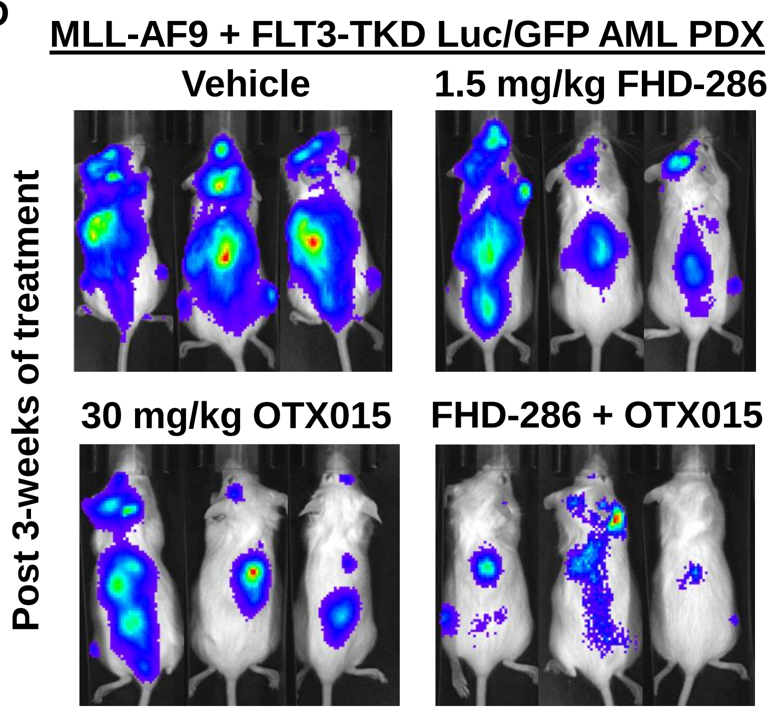

Table S1

#### Co-mutations in the MLL-AF9 PDX and their VAF

| Gene | Protein Change | Variant Allele Freq (VAF) |
|---|---|---|
| FLT3 | N676T | 0.917355 |
| KMT2C | C988F | 0.363392 |
| KMT2C | Y987H | 0.18362 |
| KMT2C | D348N | 0.078125 |
| KMT2C | V919L | 0.073446 |
| KMT2C | S990G | 0.07 |
| KMT2C | G315S | 0.04148 |
| KMT2C | R380L | 0.041427 |
| KMT2C | R284Q | 0.035762 |
| KMT2C | C391* | 0.033722 |
| KMT2C | P309S | 0.031927 |
| KMT2D | P2107T | 0.521739 |
| NOTCH2 | G225R | 0.395604 |
| NOTCH2 | C19W | 0.448276 |
| ARID1A | L1724R | 0.448276 |
| IRF8 | *427Yext*24 | 0.540881 |
| ABCA7 | P1503L | 0.539519 |
| VPS13A | S734N | 0.620079 |

#### Co-mutations identified in mtNPM1, FLT3-ITD + FLT3 D835Y AML PDX

| Gene | Protein Change | Variant Allele Freq (VAF) |
|---|---|---|
| FLT3 | ITD | 0.50 |
| FLT3 | D835Y | 0.43 |
| NPM1 | W288Cfs*12 | 0.381 |
| PHF6 | G275E | 0.940594 |
| ATRX | C1576F | 0.980583 |

**FLT3-ITD: start position chr13:28608259 (51-bp)**
**AGCCAGCTACAGATGGTACAGGTGACCGGCT**
**CCTCAGATAATGAGTACTCC**
