## Supplemental Materials and Methods for "BRG1/BRM inhibitor targets AML stem cells and exerts superior preclinical efficacy combined with BET or Menin inhibitor"

**Supplemental Methods:**

**Contact for Reagent sharing. Kapil N Bhalla. Department of Leukemia, MD. Anderson Cancer Center, 1400 Holcombe Blvd, Unit428, Houston, TX, 77030.**

**Reagents and antibodies.** FHD-286 was obtained under a material transfer agreement with Foghorn Therapeutics (Cambridge, MA). Venetoclax, OTX015, decitabine, SNDX-50469 and SNDX-5613 for in vivo studies were obtained from MedChem Express (Monmouth Junction, NJ). All compounds for in vitro studies were prepared as 10 mM stocks in 100% DMSO and frozen at -80°C in 5-10 µL aliquots to allow for single use, thus avoiding multiple freeze-thaw cycles that could result in compound decomposition and loss of activity. Anti-c-Myc [RRID: AB_1903938], anti-PUMA [RRID:AB_2797920], and anti-p21 Waf1/Cip1 [RRID:AB_823586] antibodies were obtained from Cell Signaling Technologies (Beverly, MA). Anti-BRG1 [Cat# ab110641, RRID:AB_10861578], anti-FLT3 [Cat# ab245116] and anti-CD11b [RRID:AB_2650514] antibodies were obtained from Abcam (Cambridge, MA). Anti-BCL2 [RRID: AB_626733], and anti-GAPDH [RRID: AB_627679] antibodies were obtained from Santa Cruz Biotechnologies (Dallas, TX). Anti-p27 [RRID:AB_397636] and anti-p53 [RRID:AB_395348] antibodies were obtained from BD Biosciences (Franklin Lakes, NJ). Anti-H3K27Ac [RRID:AB_2793305] antibody was obtained from Active Motif (Carlsbad, CA).

**Cell lines and cell culture.** MOLM13 [DSMZ Cat# ACC-554, RRID: CVCL_2119] and OCI-AML3 [DSMZ Cat# ACC-582, RRID:CVCL_1844] cells were obtained from the DSMZ (Braunschweig, Germany). MV4-11 [ATCC Cat# CRL-9591, RRID:CVCL_0064] cells were obtained from the ATCC (Manassas, VA). MOLM13 cells with isogenic TP53 mutations [R175H and R248Q] were a gift from Dr. Benjamin L. Ebert (Dana Farber Cancer Center, Boston, MA). HEK-293T [RRID:CVCL_0063] cells were obtained from the Characterized Cell Line Core Facility at M.D. Anderson Cancer Center, Houston TX. All experiments with cell lines were performed within 6 months after thawing or obtaining from ATCC or DSMZ. MOLM13 and OCI-AML3 cells were cultured in RPMI-1640 media with 20% FBS, 1% penicillin/streptomycin and 1% non-essential amino acids. MV4-11 cells were cultured in ATCC-formulated IMDM media with 20% FBS, 1% penicillin/streptomycin and 1% non-essential amino acids. HEK-293T cells were cultured in high-glucose-formulated DMEM media with 10% FBS, 1% penicillin/streptomycin and 1% glutamine. Logarithmically growing, mycoplasma-negative cells were utilized for all experiments. Following drug treatments, cells were washed free of the drug(s) prior to the performance of the studies described. MV4-11-MITR and OCI-AML3-MITR cells were generated by culturing MV4-11 or OCI-AML3 cells in their $LD_{90}$ concentration of FHD-286 for 96 hours. Dead cells were removed by Ficoll

Hypaque centrifugation. Live cells were washed once with complete media to remove residual Ficoll and cultured in complete media until viability was greater than 90% by trypan blue dye exclusion assessment. This process was repeated for a total of 10 shocks.

**Whole Exome Analysis of MV4-11, MV4-11-MITR, OCI-AML3 and OCI-AML3-MITR cells.** Whole exome analysis was performed on MV4-11, MV4-11-MITR, OCI-AML3 and OCI-AML3-MITR cells utilizing Agilent Exome 7 (SureSelect Human All Exon v7). The raw paired-end (PE) reads in FASTQ format were aligned to the human reference genome (hg38) for human DNA-Seq, using BWA alignment software.

**Cell Line Authentication**. The cell lines utilized in these studies were authenticated in the Characterized Cell Line Core Facility at M.D. Anderson Cancer Center, Houston TX utilizing STR profiling.

**Primary AML blasts:** Patient-derived AML cells samples were obtained with informed consent as part of a clinical protocol approved by the Institutional Review Board of The University of Texas, M.D. Anderson Cancer Center. Normal hematopoietic progenitor cells (HPCs) were obtained from delinked, de-identified cord blood samples. Mononuclear cells were purified by Ficoll Hypaque (Axis Shield, Oslo, Norway) density centrifugation following the manufacturer's protocol. Mononuclear cells were washed once with sterile 1X PBS and suspended in complete RPMI media containing 20% FBS and counted to determine the number of cells isolated prior to immuno-magnetic selection. CD34+ AML blast progenitor cells were purified by immuno-magnetic beads conjugated with anti-CD34 antibody following the manufacturer's protocol (StemCell Technologies, Vancouver, British Columbia) prior to utilization in the cell viability assays, RNA expression, and immunoblot analyses.

**Sequencing of primary de novo AML blast cells:** We performed targeted next-generation sequencing (NGS) of DNA samples from bone marrow or peripheral blood collected from patients at our center with de novo AML (**1**). Diagnostic bone marrow samples were obtained for mutational analysis. Total genomic DNA was extracted from unenriched peripheral blood (PB) or bone marrow (BM) samples using ReliaPrep genomic DNA isolation kit (Promega Corp, Madison, WI, USA). Briefly, a total of 250 ng of DNA was utilized to prepare sequencing libraries using Agilent HaloPlex custom Kit (Agilent Technologies, Santa Clara, CA, USA). The entire coding sequences of 81 genes including ABL1, ASXL1, BRAF, CALR, DNMT3A, EGFR, EZH2, FLT3, GATA1, GATA2, HRAS, IDH1, IDH2, KIT, KRAS, MDM2, IKZF2, JAK1, JAK2, MLL, MPL, MYD88, NOTCH1, NF1, NPM1, NRAS, PTPN11, RUNX1, TET2, TP53, and WT1 were interrogated on a custom-designed next-generation sequencing approach using the Illumina MiSeq platform (Illumina; San Diego, CA, USA; RRID:SCR_016379). The genomic reference sequence used was genome GRch37/hg19.

The following software tools were utilized in the experimental setup and data analysis: Illumina Experiment Manager 1.6.0 (Illumina; San Diego, CA, USA), MiSeq Control Software 2.4 (Illumina; San Diego, CA, USA), Real Time Analysis 1.18.54 (Illumina; San Diego, CA, USA), Sequence Analysis Viewer 1.8.37 (Illumina; San Diego, CA, USA), MiSeq Reporter 2.5.1 (Illumina; San Diego, CA, USA), and SureCall 3.0.1.4 (Agilent Technologies; Santa Clara, CA, USA). A minimum of 80% reads at quality scores of AQ30 or higher were required to pass quality control. The lower limit of detection of this assay (analytical sensitivity) for single nucleotide variations was determined to be 5% (one mutant allele in the background of nineteen wild type alleles) to 10% (one mutant allele in the background of nine wild type alleles). Testing of patients with active hematologic malignancies was limited to somatic mutations only.

**ChIP-Seq analysis of epigenetic state in AML cells *in vitro***. We also determined the H3K27Ac status and BRG1 occupancy in untreated and FHD-286-treated MOLM13 cells by ChIPmentation following a previously described protocol (**2**). ChIP-Seq libraries were generated with a Nextera DNA Library Preparation Kit containing the mutant Tn5 transposase (Illumina, San Diego, CA; Catalog number: FC-121-1030). The DNA fragments were indexed utilizing a Nextera Index Kit (Illumina, San Diego, CA; Catalog number: FC-121-1011) and amplified by PCR utilizing NEBNext® High-Fidelity 2X PCR Master Mix according to the manufacturer's protocol (New England Biolabs, Ipswich, MA). Library fragments were amplified for 12-15 cycles utilizing the denaturation, annealing, and extension times as previously described (**2**). The amplified library fragments were PCR-purified with a Qiagen MinElute column (Qiagen, Germantown, MD) then size selected with a 0.65X bead volume to remove large fragments (remaining on the beads), then the supernatant was combined with a 1.0X SPRI bead volume to remove fragments shorter than 200 bp. Library fragments were incubated with AMPure XP SPRI beads (Beckman Coulter, Indianapolis, IN) for 10 minutes at room temperature in 1.5 mL microcentrifuge tubes. The mixture was placed on a magnetic stand for 10 minutes. The supernatant was removed and the SPRI beads were washed twice with fresh 80% ethanol (30 seconds each wash) and air-dried for 2-3 minutes. Library DNA was eluted from the SPRI beads with a 20 µL volume of 10 mM Tris-HCl (pH 8.5). Beads were incubated at room temperature for 10 minutes, then the tubes were transferred to a magnetic stand for 10 minutes. The supernatant containing the DNA libraries was carefully removed by pipetting and transferred into a clean microcentrifuge tube. The individual libraries were quantified by Thermo Fisher Qubit [Thermo Fisher Qubit fluorimeter, RRID:SCR_018095] fluorometric quantification and quality checked by Agilent Bioanalyzer 2100 [Agilent 2100 Bioanalyzer Instrument, RRID:SCR_019389] analysis, respectively. Individual libraries were pooled into one tube, purified over a Qiagen MinElute column [QIAGEN, RRID:SCR_008539], eluted in 20 µL of 10 mM Tris, (pH 8.5) and sequenced on a NextSeq 500 next generation sequencer (Illumina

NextSeq 500, RRID:SCR_014983) utilizing a 150 cycle high-output kit (Illumina, San Diego, CA). Raw sequencing data was mapped using TopHat2 [TopHat, RRID:SCR_013035] onto the human genome build UCSC hg38 (NCBI 51) for human data [UCSC Genome Browser, RRID:SCR_005780]. Log2 fold-change in peak densities were calculated with diffReps (**3**) [diffReps, RRID:SCR_010873]. Sequence tracks were visualized with IGV software (**4, 5**) [RRID:SCR_011793]. To identify super enhancers, we performed a ranked order of super enhancers (ROSE) analysis [ROSE, RRID:SCR_017390] utilizing the H3K27Ac status of the chromatin according to the methods of Loven et al (**6**). Analysis of transcription factor binding motifs in lost BRG1 or H3K27Ac peaks was performed with HOMER [HOMER, RRID:SCR_010881].

**Bulk ATAC-Seq analysis**. ATAC-Seq libraries were generated with a Nextera DNA Library Preparation Kit containing the mutant Tn5 transposase (Illumina, San Diego, CA; Catalog number: FC-121-1030). The DNA fragments were indexed utilizing a Nextera Index Kit (Illumina, San Diego, CA; Catalog number: FC-121-1011) and amplified by PCR utilizing NEBNext® High-Fidelity 2X PCR Master Mix according to the manufacturer's protocol (New England Biolabs, Ipswich, MA). Library fragments were amplified for 10-11 cycles utilizing standard denaturation, annealing, and extension times. The amplified library fragments were PCR-purified with a Qiagen MinElute column (Qiagen, Germantown, MD) then size selected with a 1.0X bead concentration to remove fragments shorter than 200 bp. Library fragments were incubated with AMPure XP SPRI beads (Beckman Coulter, Indianapolis, IN) for 10 minutes at room temperature in 1.5 mL microcentrifuge tubes. The mixture was placed on a magnetic stand for 10 minutes. The supernatant was removed and the SPRI beads were washed twice with fresh 80% ethanol (30 seconds for each wash) and air-dried for 2-3 minutes. Library DNA was eluted from the SPRI beads with a 20 µL volume of 10 mM Tris-HCl (pH 8.5). Beads were incubated at room temperature for 10 minutes, then the tubes were transferred to a magnetic stand for 10 minutes. The supernatant containing the DNA libraries was carefully removed by pipetting and transferred into a clean microcentrifuge tube. The individual libraries were quantified and quality-checked by Thermo Fisher Qubit [Thermo Fisher Qubit fluorimeter, RRID: SCR_018095] fluorometric quantification and Agilent Bioanalyzer 2100 [Agilent 2100 Bioanalyzer Instrument, RRID: SCR_019389] analysis, respectively. Individual libraries were pooled into one tube, purified over a Qiagen MinElute column [QIAGEN, RRID:SCR_008539], eluted in 20 µL of 10 mM Tris, (pH 8.5) and sequenced on a NextSeq 500 next generation sequencer (Illumina NextSeq 500, RRID:SCR_014983) utilizing a 150 cycle mid-output kit (Illumina, San Diego, CA). Raw sequencing data was mapped using TopHat2 (**7**) [TopHat, RRID: SCR_013035] onto the human genome build UCSC hg38 (NCBI 51) for human data. Log2 fold-changes in differential ATAC-Seq peaks were calculated with diffReps (**3**) [diffReps, RRID: SCR_010873]. Sequence tracks were visualized with IGV software (**4, 5**) [RRID: SCR_011793].

**Transcriptome Analysis.** Total RNA was isolated from MOLM13 cells treated with 100 nM of FHD-286 for 16 hours utilizing a PureLink RNA Mini kit from Ambion, Inc. following the manufacturer's protocol. (Austin, TX). Sequencing libraries were prepared with ERCC spike-in controls in the MD Anderson Cancer Center DNA Sequencing and Microarray core facility and sequenced on an Illumina HiSeq-4000 next generation sequencer [Illumina HiSeq 3000/HiSeq 4000 System, RRID:SCR_016386]. Each library yielded 30-40 million read pairs. Data was mapped using STAR and Samtools (STAR [RRID: SCR_004463], SAMTOOLS [SAMTOOLS, RRID: SCR_002105]) (**8, 9**) onto the human genome build UCSC hg38 (NCBI 51) for human data. Gene expression was assessed using DESeq2 (**10**) [DESeq2, RRID: SCR_015687], then variance stabilization and quantile normalization were applied. We considered that significance was achieved for fold changes greater than or equal to 1.25X up or down relative to the untreated or parental cells, and p-values less than 0.05. The final p-values were adjusted using the Benjamini & Hochberg method (**11**). We inferred enriched pathways using the Gene Set Enrichment (GSEA) method (Gene Set Enrichment Analysis, RRID: SCR_003199) (**12**), and the gene set collection from the Molecular Signature Database (MSigDB) (**13**) [Molecular Signatures Database, RRID:SCR_016863]. Concordance between ATAC and RNA-Seq data were determined using standard bioinformatics pipelines (**14**). Circos plots were generated following the methods of Krzywinski et al. 2009 (**15**).

**TaqMan assays for detection of Menin hotspot mutations in MV4-11-MITR and OCI-AML3-MITR cells**. We designed custom TaqMan single nucleotide polymorphism probes to detect the reported hotspot mutations S160T, M327V, M327I, G331D, G331R, and T349M in the Menin gene. Twenty nanograms of genomic DNA were utilized for each assay with a TaqMan Genotyping Master Mix (Catalog # 4371353) following the manufacturer's recommended protocol and qPCR thermocycler conditions. The differences in the detection of the wild-type Menin residue versus the mutant residue is shown as Delta RN.

**Assessment of leukemia cell differentiation.** Following treatment with FHD-286, cells were harvested and washed with 1X PBS. Cells were re-suspended in 0.5% BSA/PBS and stained with APC-conjugated anti-CD11b antibody [RRID: AB_398456] or APC-conjugated IgG1 isotype control antibody [RRID: AB_398613] in the dark, at 4°C for 15-20 minutes. Cells were washed with 0.5% BSA/PBS by centrifugation at 125 x g for 5 minutes, and then re-suspended in 0.5% BSA/PBS for analysis by flow cytometry. Cells were assessed in the FL-4 fluorescence channels on a BD Accuri CFLow6 flow cytometer. Differentiation of leukemia cells was also determined by examination of cellular/nuclear morphology. Cells were cytospun onto glass slides at 500 rpm for 5 minutes. The cytospun cells were fixed and stained with a Protocol® HEMA3 stain set (Fisher Scientific, Kalamazoo, MI). Cellular/nuclear morphology was assessed by light microscopy. Two hundred cells were counted in at least 5 different sections of the slide for each condition. The % morphologic differentiation is reported relative to control cells. Each experiment was performed at least twice.

**Assessment of percentage non-viable cells.** Following designated treatments (72-96 hours), cultured cell lines or patient-derived (PD) AML blast cells, were washed with 1X PBS, stained with TO-PRO-3 iodide (Cat# T3605, Life Technologies, Carlsbad, CA) and analyzed by flow cytometry on a BD Accuri CFlow-6 flow cytometer (BD Biosciences, San Jose, CA). We used matrix dosing of agents in combinations to allow synergy assessment utilizing the SynergyFinder V2 online web application tool and Delta Synergy scores by ZIP method (http://synergyfinder.fimm.fi/) (**16-18**).

**Single cell multiomics (combined ATAC-Seq and RNA-Seq) analysis.** PD mtNPM1+FLT3-ITD AML#19, MLL-AF9 AML#14 and MLL-AF6 AML#16 cells were treated with 100 nM of FHD-286 (~2 million cells per condition) for 16 hrs and then cryopreserved in freezing media (90% FBS + 10% DMSO) until processing of the samples occurred. Nuclei preparation for combined sc-ATAC-Seq and scRNA-Seq followed the recommended demonstrated protocol from 10X Genomics. Isolated nuclei were treated with Tn5 transposase following the manufacturer's recommended protocol and then separated into gel bead emulsions utilizing the 10X Genomics Chromium Controller and Chip J kit. Pre-amplification, indexing, and library preparations were conducted following the manufactures recommendation (10X Genomics). Raw scRNA-seq data was pre-processed, de-multiplexed, and aligned to human reference genome (GRCh38) using CellRanger (10X Genomics). Cells with few (<200) or many (>6,000) genes, likely doublets or multiplets predicted by Scrublet (Wolock et al. 2019), and cells with >20% of read counts derived from the mitochondrial genome were removed. The batch effects were corrected by Harmony (**19**) and Seurat v4 (**20**). Raw unique molecular identifier (UMI) counts were log-normalized and used for principal component analysis (PCA). Seurat v4 (**20**) was applied to the normalized gene-cell matrix to identify highly variable genes for unsupervised cell clustering. For visualization, the dimensionality was further reduced using Uniform Manifold Approximation and Projection (UMAP) (**21**) method. We also sub-clustered cells within each lineage to identify transcriptomically distinct subpopulations. To define the major cell type and state of each single cell, the top 50 most significant differentially expressed genes (DEGs) were identified for each cell cluster. We also identified DEGs for cell subpopulations of interest using Seurat which was then filtered to select significant DEGs (log2 fold change >1.0 or <-1.0 and FDR p-value <0.05) between two conditions. For pathway analysis, the curated gene sets (including Hallmark, GO, KEGG, REACTOME gene sets) was downloaded from the Molecular Signature Database (**13**) [Molecular Signatures Database, RRID: SCR_016863], and single-sample GSVA was applied and pathway scores was calculated for each cell type using the GSVA software package (**22**). Gene set enrichment analysis using GSEA software package (**12**) was performed to identify significantly enriched signaling pathways (FDR p-value < 0.05) between two conditions.

**Integrated analysis of sc-ATAC-Seq and sc-RNA-Seq data.** For paired scATAC-seq data, Signac (v1.9.0) (**23**) was used to process the fragment files. We removed low-quality cells based on peak region fragments, nucleosome signal and TSS enrichment. The accessible chromatin peaks were called by using MACS2 (**24**) method with the default parameters of Signac's CallPeaks function. After removing peaks in non-standard chromosomes and in genomic blacklist regions, we quantified counts in each peak using the FeatureMatrix function. Then, we performed latent semantic indexing (LSI) and normalized with term-frequency inverse-document-frequency (TFIDF). RunSVD function was used on the TFIDF matrix to consequent dimensionality reduction with singular value decomposition (SVD). Sc-ATAC-Seq and sc-RNA-Seq data were integrated by using Seurat. For each cell type, we found differentially accessible peaks between groups by the "FindAllMarkers" function. Based on the filtered, differentially accessible chromatin regions (log2 fold change >1.0 or <-1.0 and FDR p-value <0.05), we performed enriched motifs analysis in these genomic regions by HOMER (**25**). We used "filterbarcodes" command of sinto (v0.9.0, https://timoast.github.io/sinto/) to get bam files for interested cell type, and generated bigWig files by using "bamCoverage" command of deeptools (**26**). BigWig files were uploaded to the UCSC genome browser [UCSC Genome Browser, RRID:SCR_005780] (**27**) for visualization.

**Plasmid Generation, Viral Packaging, and Creation of Cell Lines.** Plasmid constructs for the production of lentivirus were transfected with packaging plasmids psPAX2 and pMD2.G into HEK-293T cells utilizing jetPRIME reagent (PolyPlus Transfection, New York, NY). The psPAX2 and pMD2.G packaging plasmids were a gift from Didier Trono (Addgene plasmid #12260 and #12259 [RRID: Addgene_12260; RRID: Addgene_12259]). Media was changed the following day. Viral supernatant was collected 72 hours post transfection and filtered through a 0.45 µm PES membrane. AML cells were seeded at 5 x $10^5$ cells/mL in a 50:50 mix of media and lentivirus supernatant with 8 µg/mL polybrene (Sigma-Aldrich, Burlington, MA). The following day, the viral supernatant was removed by centrifugation and cells were transduced with fresh viral supernatant for an additional 24 hours. To generate luciferase-expressing AML cells, pHIV-Luc-ZsGreen (a gift from Bryan Welm [Addgene plasmid #39196; http://n2t.net/addgene:39196; RRID:

Addgene_39196]) was packaged as above and transduced into patient-derived AML PDX cells. ZsGreen-positive cells were sorted by flow cytometry (FACSAria, FL-1 channel, top 10% brightest GFP-expressing cells), and expanded in culture or grown in NSG mice prior to their utilization in therapeutic in vivo mouse studies.

**RNA isolation and quantitative polymerase chain reaction.** Following the designated treatments, total RNA was isolated from AML cells utilizing a PureLink RNA Mini kit from Ambion, Inc. (Austin, TX) and reverse transcribed with a High Capacity Reverse Transcription kit from Life Technologies (Carlsbad, CA). Quantitative real-time PCR analysis for the expression of target genes was performed on cDNA using TaqMan probes and a TaqMan Universal PCR Master Mix from Applied Biosystems (Foster City, CA). Relative mRNA expression was normalized to the expression of GAPDH and compared to the untreated cells.

**Cell lysis and protein quantitation.** Untreated or drug-treated cells were centrifuged, and the cell pellets were incubated in lysis buffer on ice for 20 minutes (**28**). After centrifugation, an aliquot of each cell lysate was diluted 1:10 and the protein content was quantitated using a BCA protein quantitation kit (Pierce, Rockford, IL), according to the manufacturer's protocol. Protein concentrations were determined by comparing the absorbance at 562 nm compared to a known concentration range of bovine serum albumin (BSA) from 0.125 mg to 2 mg/mL.

**SDS-PAGE and immunoblot analyses.** Thirty micrograms of total cell lysate were used for SDS-PAGE. Western blot analyses were performed on total cell lysates using specific antisera or monoclonal antibodies. Blots were washed with 1X PBST, then incubated in IRDye 680RD goat anti-mouse (RRID: AB_10956588) or IRDye 800CW goat anti-rabbit (RRID: AB_621843) secondary antibodies (LI-COR, Lincoln, NE) for 1 h, washed three times in 1X Phosphate Buffered Saline with Tween®20 (PBST) and scanned with an Odyssey CLX Infrared Imaging System utilizing Image Studio 5.0 Software (RRID: SCR_015795) (LI-COR, Lincoln, NE). The expression levels of β-Actin or GAPDH in the cell lysates were used as the loading control for the western blots. Immunoblot analyses were performed at least twice.

**Single cell next-generation mass cytometry 'CyTOF' analysis of MLL1r and mtNPM1-expressing AML cells.** Primary, patient-derived MLL1r and mtNPM1-expressing AML cells were treated with 100 nM of FHD-286 for 48 hours. At the end of treatment, cells were blocked with staining buffer (0.5% BSA/PBS) for 30 minutes, then a cocktail of extracellular antibodies (CLEC12A, CD123, CD244, CD99, CD33 and CD11b) conjugated to transition element isotopes were added and incubated for 1 hour at room temperature (RT). For viability staining, a 5 µM concentration of cisplatin was added and incubated at RT for 2 minutes. Cells were washed with staining buffer, centrifuged at 500 x g for 5 minutes and staining buffer was vacuum aspirated. Cells were fixed with 100 µL of 1.6% paraformaldehyde (PFA) for 10 minutes at room temperature. Following this, cells were permeabilized with 900 µL of ice-cold 100% methanol (90% volume) at -20°C for at least 20 minutes. Next, cells were washed with 1 ml of staining buffer to remove the paraformaldehyde/methanol solution. Cells were blocked in 50 µL of staining buffer for 30 minutes and a cocktail of intracellular antibodies conjugated to transition element isotopes was added to be used as tags in atomic mass spectrometric analysis of the cells. Cells were incubated for 1 hour at room temperature, then washed with staining buffer at 500 x g for 5 minutes. Intercalator was added (500 µL of 1:1000 Ir-intercalator diluted in 1.6% PFA/1X PBS) and cells were incubated overnight at 4°C. Cells were washed twice (500 x g for 5 minutes per wash) in staining buffer, then counted using a Countess II counting device. Following the last wash, $1 \times 10^6$ cells were suspended in 100 µL of de-ionized water and incubated overnight at 4°C. Time-of-flight mass spectrometry (CyTOF) measured multiple different cellular parameters simultaneously in each cell. The absolute fold-change of protein expression changes in FHD-286-treated cells over control cells within the CLEC12A Hi, CD123 Hi, CD99 Hi, CD33 Hi, CD11b Lo population was analyzed by the Astrolabe Cytometry Platform (Astrolabe, Fort Lee, NJ).

**Proteome Profiling.** Protein extraction, digestion and peptide fractionation was carried out based on the protocol adapted from (**29**). Briefly cells were lysed in 8M urea buffer, reduced/alkylated and digested using LysC and Trypsin proteases. The peptides from MOLM13 cells and DF16835 cells were labeled with TMT10 and TMTpro 16-plex isobaric label reagent (Thermo Fisher Scientific) respectively according to manufacturer's protocol. The high-pH offline fractionation was carried to generate 24 peptide pools. The deep-fractionated peptide samples were separated on an online nanoflow Easy-nLC-1200 system (Thermo Fisher Scientific) and analyzed on Orbitrap Exploris 480 mass spectrometer (Thermo Fisher Scientific). 1 µg of each fraction was loaded on a pre-column (2 cm x 100 µm I.D.) and separated on in-line 20 cm x 75 µm I.D. column (Reprosil-Pur Basic C18 aq, Dr. Maisch GmbH, Germany) equilibrated in 0.1% formic acid (FA). Peptide separation was done at a flow rate of 200 nl/min over 110 min gradient time with different concentration of 90% acetonitrile solvent B (2% to 30% 87 min, 30% to 60% 6 min, 60% to 90% 7 min and finally hold at 50% 10 min). The heated column was maintained at 60°C. The mass spectrometer was operated in a data dependent mode with 2 second cycle time. The MS1 was done in Orbitrap (120000 resolution, scan range 375-1500 m/z, 50 ms injection time) followed by MS2 in Orbitrap at 30000 resolution (HCD 38%) with TurboTMT algorithm. Dynamic exclusion was set to 20sec and the isolation width was set to 0.7m/z. The mass spectra were searched using MSFragger (v3.5). The reverse decoys and common contaminants were added to the NCBI Refseq human protein database (downloaded 2021_12_23) using Philosopher (**30**). The raw data processing and differential analysis was carried out as described before (**31**). Gene Set Enrichment Analysis (GSEA) (**12**) was performed using the canonical pathway gene sets derived from KEGG, REACTOME and HALLMARK gene sets (**13**) and oncogenic signature gene sets.

**In vivo FHD-286 treatment in immune-competent mice**. C57BL/6J mice (Stock #: 000664, 6 weeks of age) [Jackson Labs, Bar Harbor, ME; RRID: IMSR_JAX: 000664], were treated with vehicle or 1.5 mg/kg of FHD-286 (n=5 mice per cohort) for two weeks. Weights of the mice were monitored on a weekly basis to adjust drug dose. Peripheral blood (50-100 µL) was collected by retro-orbital bleed under anesthesia (isoflurane) and complete blood count (CBC) analyses were performed. Following this, the mice were allowed to recover for two weeks with no treatment and CBCs were repeated.

**In vivo efficacy of FHD-286 in targeting AML-initiating stem cells.** For the ex vivo treated engraftment model, female NOD.Cg-Prkdc$^{scid}$ Il2rg$^{tm1Wjl}$/SzJ (NSG) mice (stock number: 005557; 4-6 weeks of age) [Jackson Labs, Bar Harbor, ME; RRID: IMSR_JAX:005557] were exposed to 2.5 Gy of irradiation from a cesium source. The following day, mice (n=6 mice per cohort) were injected in the lateral tail vein with 2.5 x $10^6$ luciferase-GFP expressing mtNPM1 + FLT3-ITD AML PDX cells (Dana Farber PDX number: DF16835) (**32**) that had been treated ex vivo with 0, 10 or 30 nM of FHD-286 for 96 hours. Mice were monitored daily for 4-5 days post cell infusion. Mice were imaged utilizing a Xenogen IVIS Lumina in vivo imaging system to document engraftment one week, two weeks and 4 weeks post cell infusion. Total bioluminescence was recorded as photons/second. Mice that became moribund or experienced hind limb paralysis were euthanized according to the approved IACUC protocol. Department of Veterinary Medicine staff members assisting in determining when euthanasia was required were blinded to the experimental conditions of the study. The survival of the mice is represented by a Kaplan-Meier plot. Significance was determined by a Mantel-Cox log rank test. P-values of less than 0.05 were assigned significance. To determine the effects of FHD-286 treatment on leukemia-initiating stem cells in vivo, we infused pre-irradiated (2.5Gy) NSG mice with 2.5 x $10^6$ luciferase-GFP expressing mtNPM1 + FLT3-ITD AML PDX cells and the mice were monitored daily for 4-5 days post cell infusion. Mice were imaged utilizing a Xenogen IVIS Lumina in vivo imaging system to document leukemia engraftment. Following this, mice were randomized into groups based on equivalent mean bioluminescent intensity to control for variation in cell engraftment and variation between different treatment groups. Drug treatments were initiated on day 7. Mice were treated with Vehicle or FHD-286 (1.5 mg/kg, daily x 5 days, by oral gavage) for 5 weeks. Mice were imaged weekly by bioluminescent imaging to document treatment efficacy and/or disease progression. Total bioluminescence was recorded as photons/second. Mice that became moribund or experienced hind limb paralysis during the course of the study were euthanized according to the approved IACUC protocol. When vehicle mice had to be euthanized due to excess leukemia burden or paralysis, we also sacrificed two FHD-286 treated mice and harvested the spleen and bone marrow. Human AML cells were isolated utilizing an EasySep™ Mouse/Human Chimera isolation kit (Catalog # 19849A, StemCell Technologies, Vancouver, BC). Equal numbers (2.5 x $10^6$ cells) of viable, human sorted, luciferase-GFP expressing mtNPM1 + FLT3-ITD AML PDX cells from vehicle or FHD-286 treated mice were re-transplanted into pre-irradiated NSG mice (N=6 per cohort) and the mice were monitored daily for 4-5 days post cell infusion. Mice were imaged weekly by bioluminescent imaging to document disease progression. Total bioluminescence was recorded as photons/second. During the course of the study, mice that became moribund or experienced hind limb paralysis were euthanized according to the approved IACUC protocol. Additionally, Department of Veterinary Medicine staff members assisting in determining when euthanasia was required were blinded to the experimental conditions of the study. The survival of the mice is represented by a Kaplan-Meier plot. Significance was determined by a Mantel-Cox log rank test. P-values of less than 0.05 were assigned significance.

**In vivo models of de novo AML treated with FHD-286 alone or FHD-286 based combinations.** All in vivo studies were approved by and conducted in accordance with the guidelines of the IACUC at the M.D. Anderson Cancer Center, an AAALAC-accredited facility. Male and female NOD.Cg-Prkdc$^{scid}$ Il2rg$^{tm1Wjl}$/SzJ (NSG) mice (stock number: 005557; 4-6 weeks of age) [Jackson Labs, Bar Harbor, ME; RRID: IMSR_JAX:005557] were exposed to 2.5 Gy of radiation. The following day, mice (n=4-6 mice per cohort for single agent FHD-286 leukemia models; n=7-8 per cohort for combination studies) were injected in the lateral tail vein with 2.5-3.0 x $10^6$ luciferase-GFP expressing AML PDX cells (Dana Farber PDX number: DF68555, DF16835, or PDX AML#25 from the oncoplot [mtNPM1, FLT3-ITD + FLT3-F691L]) (**32**) and monitored daily for 4-5 days. Mice were imaged utilizing a Xenogen IVIS Lumina in vivo imaging system to document engraftment before treatment was initiated. Mice were randomized into groups based on equivalent mean bioluminescent intensity to control for variation in cell engraftment and variation between different treatment groups. Treatments were initiated on day 7. For the MLL-AF9 + FLT3-TKD Luc/GFP AML PDX model, mice were treated with Vehicle, FHD-286 (1.5 mg/kg, daily x 5 days, by oral gavage), venetoclax (30 mg/kg, daily x 5 days, by oral gavage), decitabine (1 mg/kg, daily x 5 days [first week only], by subcutaneous injection), OTX015 (30 mg/kg, daily x 5 days, by oral gavage) and combinations of FHD-286 with venetoclax, decitabine or OTX015 for 4-7 weeks. For the mtNPM1 + FLT3-ITD Luc/GFP AML PDX model, mice were treated with Vehicle, FHD-286 (1.5 mg/kg, daily x 5 days, by oral gavage), OTX015 (30 mg/kg, daily x 5 days, by oral gavage), SNDX-5613 (50 mg/kg B. I. D. x 5 days, by oral gavage) and combinations of FHD-286 with OTX015 or

SNDX-5613 for 8 weeks. FHD-286 was prepared in a solution of 10% (vol/vol) DMSO followed by 90% (vol/vol) Captisol (20% solution). Decitabine was prepared in sterile 1X PBS. Venetoclax and OTX015 were prepared in a solution of 10% (vol/vol) of 95% ethanol, followed by 30% (vol/vol) of PEG-400 (ThermoFisher Scientific, Waltham, MA), and then 60% (vol/vol) of Phosal-50 (LIPOID, LLC via ThermoFisher Scientific, Waltham, MA). SNDX-5613 was prepared in a solution of 10% (vol/vol) DMSO followed by 90% (vol/vol) Captisol (20% solution). Mice were imaged weekly by bioluminescent imaging to document treatment efficacy and/or disease progression. Total bioluminescent flux was recorded as photons/second. Mice that became moribund or experienced hind limb paralysis were euthanized according to the approved IACUC protocol. Department of Veterinary Medicine staff members assisting in determining when euthanasia was required were blinded to the experimental conditions of the study. The survival of the mice is represented by a Kaplan-Meier plot. Significance was determined by a Mantel-Cox log rank test. P-values of less than 0.05 were assigned significance.

**Power analysis for in vivo studies**. With a sample size of 10 mice per group, we can achieve 79.5% power to detect a difference of overall survival at a significance level of 0.05 with one-sided log-rank test, assuming 30% of mouse-survival at the end of study in the experimental group.

**Statistical analysis**. Significant differences between values obtained in AML cells treated with different experimental conditions compared to untreated control cells were determined using the Student's t-test in GraphPad V9 [RRID:SCR_002798]. For the *in vivo* mouse models, a two-tailed, unpaired t-test was utilized for comparing total bioluminescent flux differences between vehicle and single agent treated or between single agent and combination treated mice. For survival analysis, a Kaplan-Meier plot and a Mantel–Cox log rank test were utilized for comparisons of different cohorts. P-values of < 0.05 were assigned significance.

**Data and Software availability**. ATAC-Seq, sc-ATAC-Seq, ChIP-Seq, bulk RNA-Seq and sc-RNA-Seq datasets have been deposited in GEO under Accession IDs. The mass spectrometry proteomics data have been deposited to the ProteomeXchange Consortium via the PRIDE partner repository with the dataset identifier "PXDxxxxxx"
